# VAMP8 function reveals tight linkage between endocytic recycling and endocytosis

**DOI:** 10.1101/2025.08.31.673318

**Authors:** Ailing Liu, Yueping Li, Zheng Huang, Wen Chen, Peiliu Xu, Xiangying Wei, Guosheng Hu, Shuangquan Liu, Xiaoxia Liu, Yaohui He, Danling Wang, Sandra L. Schmid, Zhiming Chen

## Abstract

Clathrin-mediated endocytosis (CME) is a multistage process that involves the initiation and stabilization of clathrin-coated pits (CCPs) that invaginate and finally detach from the plasma membrane to form clathrin-coated vesicles (CCVs). Given that SNARE proteins are essential for downstream vesicle targeting and fusion events, their recruitment into nascent CCVs has been suggested to be a prerequisite for CME progression. However, which and how SNARE proteins regulate CME remains to be explored. Here, we showed that siRNA-mediated knockdown of the R-SNARE, VAMP8 impairs CCP initiation, stabilization and invagination and strongly inhibits CME. Mechanistically, recruitment of VAMP8 to CCVs is not required for CME. Instead, depletion of VAMP8 inhibits recycling of endocytic cargoes and as exemplified here by transferrin receptor, skews their trafficking toward lysosomal degradation. VAMP8 depletion therefore indirectly impairs CCV formation and inhibits CME by depleting endocytic cargo. Overall, our study provides new insights into the crosstalk between endocytosis and endocytic recycling of CME cargo and demonstrates the critical role for cargo recruitment in stabilizing nascent CCPs to regulate CME.

**Significance Statement:** We demonstrate that VAMP8 depletion inhibits recycling of endocytic cargo and reroutes transferrin receptors toward lysosomal degradation, thereby impairing endocytic vesicle formation and inhibiting clathrin-mediated endocytosis. Thus, the work uncovers a critical link between endocytic recycling and endocytosis, highlighting the importance of recycling pathways in maintaining membrane dynamics. However, the recruitment of VAMP8 into nascent clathrin-coated pits by CALM is not required for CME. While SNARE recruitment is essential for formation of COPI and COPII vesicles, our findings suggest that this requirement is not generalizable to endocytic CCVs.

## Introduction

Clathrin-mediated endocytosis (CME) is a major endocytic pathway that regulates the uptake of receptor proteins from the plasma membrane (PM), and is thus essential for signaling, cell-cell communication, and cellular homeostasis (1, 2). Productive CME events involve a multistage process that includes the initiation and stabilization of nascent clathrin-coated pits (CCPs). These pits invaginate, mature and eventually detach from the PM to form clathrin-coated vesicles (CCVs) that are loaded with cargo (2, 3). Not all CME initiation events complete this multistage process: many nascent clathrin assemblies rapidly disassemble as early abortive coats (ACs); those that are stabilized are designated as *bona fide* CCPs, herein referred to simply as CCPs. A second point of failure occurs when CCPs fail to grow and invaginate. These are turned over as late abortive pits (4–9). The early stages of CME, including the initiation, stabilization and invagination of CCPs, are key steps in determining the fate of nascent CCPs. However, the underlying mechanisms determining their fates, and thereby the efficiency of CME, remain incompletely understood.

Soluble N-ethylmaleimide-sensitive factor attachment receptors (SNAREs) are small membrane-anchored proteins that function in complexes consisting of one R-SNARE and three Q-SNAREs, to provide the specificity and energy for targeting and membrane fusion of transport vesicles (10–13). SNARE proteins participate in almost all membrane fusion processes (12, 14), including the targeting and fusion of nascent CCVs required for delivery of their cargo to early endosomes (15).The incorporation of SNARE proteins into nascent vesicles has been shown to be required in yeast for COPI (16) and COPII (17, 18) vesicle formation, but this has not been shown for CME. However, the existence of SNARE-specific adaptor proteins, CALM (Clathrin assembly lymphoid myeloid leukemia), for VAMPs 2, 3,7 and 8 (19–21) and Hrb (HIV-1 Rev-binding protein) for VAMP 7 (22, 23) has been suggested as a mechanism to ensure incorporation of SNARE proteins into CCVs without competition from other cargo. While depletion of either Hrb or CALM inhibits CME, these proteins have other partners and are likely to play additional critical roles in CME. Indeed, point mutations in CALM that selectively inhibit SNARE uptake, uncouple their effects on SNARE uptake from CALM’s other roles in CME (24). A more recent study has shown that knockdown of several SNARE proteins alters early CCP dynamics (25), but their effects on CME were not measured.

Vesicle-associated membrane protein 8 (VAMP8, also known as endobrevin) is an R-SNARE protein that is widely distributed on the PM, early/recycling/late endosomes, and lysosomes (26–29). Best described are its involvement in regulated exocytosis (30–33), fusion of recycling endosomes with the PM (34), lysosome-autophagosome fusion (29, 35–37) and homotypic fusion of early and late endosomes (26, 38, 39). This latter function, together with its localization to CCPs and early endosomes (26) and an earlier finding that VAMP8 knockdown, in the context of a CRISPRi-based screen impairs CCP initiation and stabilization (25), led us to further characterize the role of VAMP8 in CME.

In this study, we reveal that VAMP8 knockdown impairs CCP initiation, stabilization and invagination and thus inhibits CME. However, the effects on CME are not determined by the direct recruitment of VAMP8 to CCPs. Instead, VAMP8 depletion inhibits CME indirectly by mediating endocytic recycling of cargo from early and recycling endosomes to the PM. Knocking down VAMP8 skews the sorting of endosomal CME cargo, shown here for transferrin receptor (TfnR), from recycling towards lysosomal degradation. We show that the resulting reduction in CME cargo at the plasma membrane inhibits CCV formation. Overall, our study provides new insights into the crosstalk between endocytosis and exocytosis intersected by VAMP8-mediated cargo recycling and demonstrates the significance of cargo recruitment in regulating early stages of CME.

## Results

### VAMP8 knockdown inhibits CCP maturation and TfnR uptake

To determine the role of VAMP8 in CME, we first performed siRNA-mediated knockdown of VAMP8 in ARPE-HPV cells that stably express eGFP-tagged clathrin light chain a (referred to as ARPE-HPV eGFP-CLCa) and examined its effects on CCP initiation and maturation. Two sets of siRNA were used separately and both efficiently knocked down VAMP8 (Fig. 1A and B). Cells were time-lapse imaged using Total Internal Reflection Fluorescence Microscopy (TIRFM) (40, 41). Strikingly, the number of clathrin-coated structures (CCSs, which include both transient and stable clathrin-labeled structures) on the cell surface was strongly reduced after VAMP8 knockdown (Fig. 1C-E and Movie 1). To better quantify CCP dynamics and further determine which stage(s) of CME were effected by VAMP8 depletion, we analyzed the acquired imaging data with cmeAnalysis (7, 9, 42) and disassembly asymmetry score classification (DASC). Together these analytical pipelines provide a comprehensive and unbiased characterization of CCP intermediates and CME progression (5). Software analysis revealed that VAMP8 knockdown not only significantly inhibited the initiation rate of CCSs (Fig. 1F), but also the initiation rate (Fig. 1G) and percentage of CCPs (CCP%, Fig. 1H). These findings confirm our previous phenotypic analysis of VAMP8 knockdown.(25). In addition, VAMP8 knockdown extends the lifetime of CCPs (Fig. 1I) while leading to a shift toward dimmer (i.e. smaller or flatter) structures (Fig. 1J), indicating that CCP maturation was also inhibited in the absence of VAMP8. Importantly, these effects could be rescued by expression of siRNA-resistant VAMP8 (Fig. S1A-D).

**Figure 1.**
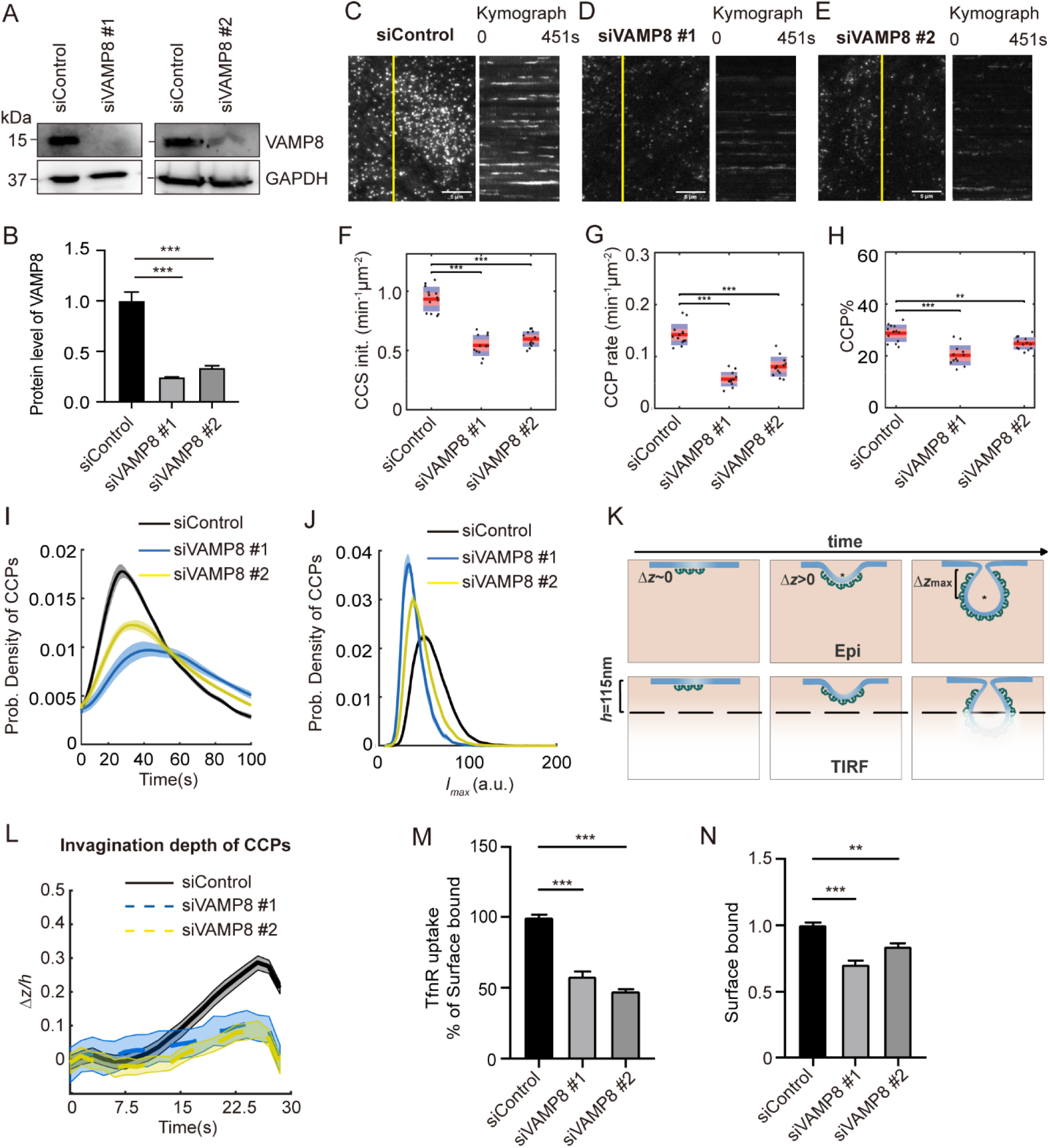
VAMP8 knockdown inhibits CCP maturation and TfnR uptake. **(A)** ARPE-HPV eGFP-CLCa cells were transfected with control or VAMP8 siRNA. Two sets of siRNAs were used to knock down VAMP8. Western blotting was used to determine the knockdown efficiency of VAMP8. **(B)** Quantification of the VAMP8 knockdown efficiency. Error bars indicate SEM of three independent experiments. **(C-E)** Representative single frame images and corresponding kymographs (for region indicated by yellow lines) from TIRFM movies (1 frame/s, 7.5 min/movie, see Movie 1) that were captured in ARPE-HPV eGFP-CLCa cells treated with (C) control siRNA or (D and E) VAMP8 siRNA. Scale bars = 5 µm. **(F-H)** Effect of VAMP8 knockdown on the initiation rates of (F) all CCSs and (G) CCPs, as well as (H) CCP%. Each dot represents a movie, and N = 12 movies for each condition. **(I and J)** Effect of VAMP8 knockdown on the (I) lifetime distribution and (J) Maximum fluorescence intensity (*I_max_*) distribution of CCPs. The data presented was acquired from a single experiment that represents three independent biological repeats. Number of dynamic tracks analyzed: 16,7117 for siControl, 11,0803 for siVAMP8 #1, and 10,6918 for siVAMP8 #2. Shadow area indicates 95% confidential interval. **(K)** Scheme of Epi-TIRF microscopy for measuring the invagination of CCPs using primary/subordinate tracking. Δz denotes the invagination depth of CCPs. □=115 nm is the evanescent depth of TIRF field. See Methods for more details. **(L)** Epi-TIRF microscopy analysis showed that VAMP8 knockdown significantly inhibited CCP invagination. Curves are the mean invagination traces of CCPs represented by Δ*z(t)/h*. Shadowed area indicates 95% confidence interval. The presented data was acquired from a single experiment that represents three independent repeats. N = 15 movies for each condition. Number of CCP tracks analyzed to obtain the Δ*z(t)/h* curves: 18,625 for siControl, 2,715 for siVAMP8 #1 and 3,911 for siVAMP8 #2. **(M and N)** Evaluation of the (M) uptake efficiency of TfnR (Internalized/Surface-bound) and (N) Surface bound TfnR by in-cell ELISA. Error bars indicate SEM of N = 16 samples. Statistical analysis of the data in (B, M and N) was performed using GraphPad Prism 8 by unpaired t-test. Statistical analysis of the data in (F-H) is the Wilcoxon Rank Sum test, \*\**P* ≤ 0.01, \*\*\**P* ≤ 0.001.

During CCP maturation, invagination of the clathrin-coat and its underlying membrane is a crucial determinant as to whether the nascent CCP will be either ‘productive’ (i.e. go on to form CCVs and internalize cargo) or abortive (5, 8, 9). To determine whether VAMP8 knockdown affects CCP invagination, we conducted Epifluorescence (Epi)-TIRF microscopy imaging to measure CCP invagination depth in live ARPE-HPV eGFP-CLCa cells (5, 6, 8). In this approach, Epi and TIRF channel signals are near-simultaneously imaged. As CCPs invaginate, the TIRF channel signal becomes dimmer due to the exponential-decay of the evanescent wave energy, while the Epi channel signal is still proportional to the number of eGFP-tagged clathrin, resulting in an increased ratio of Epi/TIRF that allows us to measure the invagination depth of CCPs (Fig. 1K). To yield the average time course of the invagination depth (Δz) of the center-of-mass of CCPs, cmeAnalysis tracked CCSs using a primary (TIRF-channel)/subordinate (Epi-channel) algorithm. DASC was used to distinguish CCPs from ACs. The Epi and TIRF fluorescence intensity traces of CCPs were then temporally aligned, averaged and logarithmically transformed to measure the acquisition of curvature over time (8). Here, we present the cohort of CCPs with lifetime average of ∼30s because they represent the invagination behavior of the most frequent tracks (Fig. 1I). Surprisingly, VAMP8 knockdown strongly inhibited CCP invagination (Fig. 1L), and this phenotype was rescued upon expression of siRNA-resistant VAMP8 (Fig. S1E). These results indicate an unexpected effect of VAMP8 depletion on the crucial invagination process of CME.

Next, we examined the effects of VAMP8 knockdown on the cellular uptake of a prototypical CME cargo protein, TfnR. Consistent with TIRFM analysis results, VAMP8 knockdown significantly reduced the uptake efficiency of TfnR (Fig. 1M). Interestingly, the level of TfnR on the cell surface was also significantly reduced (Fig. 1N). Typically, a reduction in the rate of TfnR endocytosis leads to an increase in surface TfnR, because the larger pool of intracellular TfnRs is recycled to and accumulate on the PM. These results suggested an imbalance between the endocytosis and endocytic recycling of TfnR.

In parallel, we examined the effects of siRNA-mediated knockdown of VAMP2 (aka synaptobrevin 2, a closely related R-SNARE) on CME in ARPE-HPV eGFP-CLCa cells. VAMP2 is a neuron-abundant protein that plays a crucial role in neuronal synaptic transmission but is also expressed in ARPE-HPV and other nonneuronal cell types. Interestingly, depletion of VAMP2 in ARPE-HPV cells did not affect CCP initiation, maturation or TfnR uptake (Fig. S2 and Movie 2), which further strengthens the specificity of VAMP8’s role in CME in non-neuron cells.

Together these observations suggested a critical role for VAMP8 during multiple stages of CME. Next we explored the mechanism underlying these effects on VAMP8 depletion on CME.

### Abolishing VAMP8 recruitment to CCVs does not affect CME

Given that incorporation of a sufficient number of SNARE proteins into nascent CCVs has been suggested to be critical for CCP maturation (25), we first tested whether the direct recruitment of VAMP8 into CCPs is required.

CALM has been identified as a VAMP8 adaptor protein (20). L219S/M244K mutations in the VAMP8 binding site on CALM prevent the recruitment of VAMP8 to CCPs (20). To examine the effect of abolishing VAMP8 recruitment to CME, we introduced these mutations into CALM and generated two cell lines: 1) ARPE-HPV eGFP-CLCa cells that stably express RFP-VAMP8 and siRNA-resistant, myc-tagged CALM (referred to as: CALM(WT)); 2) ARPE-HPV eGFP-CLCa cells that stably express RFP-VAMP8 and siRNA-resistant, myc-tagged CALM(L219S/M244K) (referred to as: CALM(SNARE*)) (Fig. S3). The RFP-VAMP8 construct was confirmed to be functional, as its expression rescued the CCP initiation, stabilization and curvature generation defects caused by VAMP8 knockdown (Fig. S4).

Endogenous CALM was depleted with specifically targeted siRNA (Fig. S3E) and the distribution of VAMP8 was assessed by immunofluorescence. In cells expressing CALM(WT), VAMP8 appears in small clusters on the PM, many of which colocalize with CCPs (Fig. 2A). In contrast, and consistent with a previous report (20), VAMP8 is diffusely distributed on the PM of cells expressing CALM(SNARE*) (Fig. 2B). Moreover, dual-channel time-lapse TIRFM imaging (5, 9, 41, 42) confirmed that VAMP8 can be recruited to CCPs in cells expressing CALM(WT) (Fig. 2C) but not in cells expressing CALM(SNARE*) (Fig. 2D), even though surface levels of VAMP8 are substantially increased.

**Figure 2.**
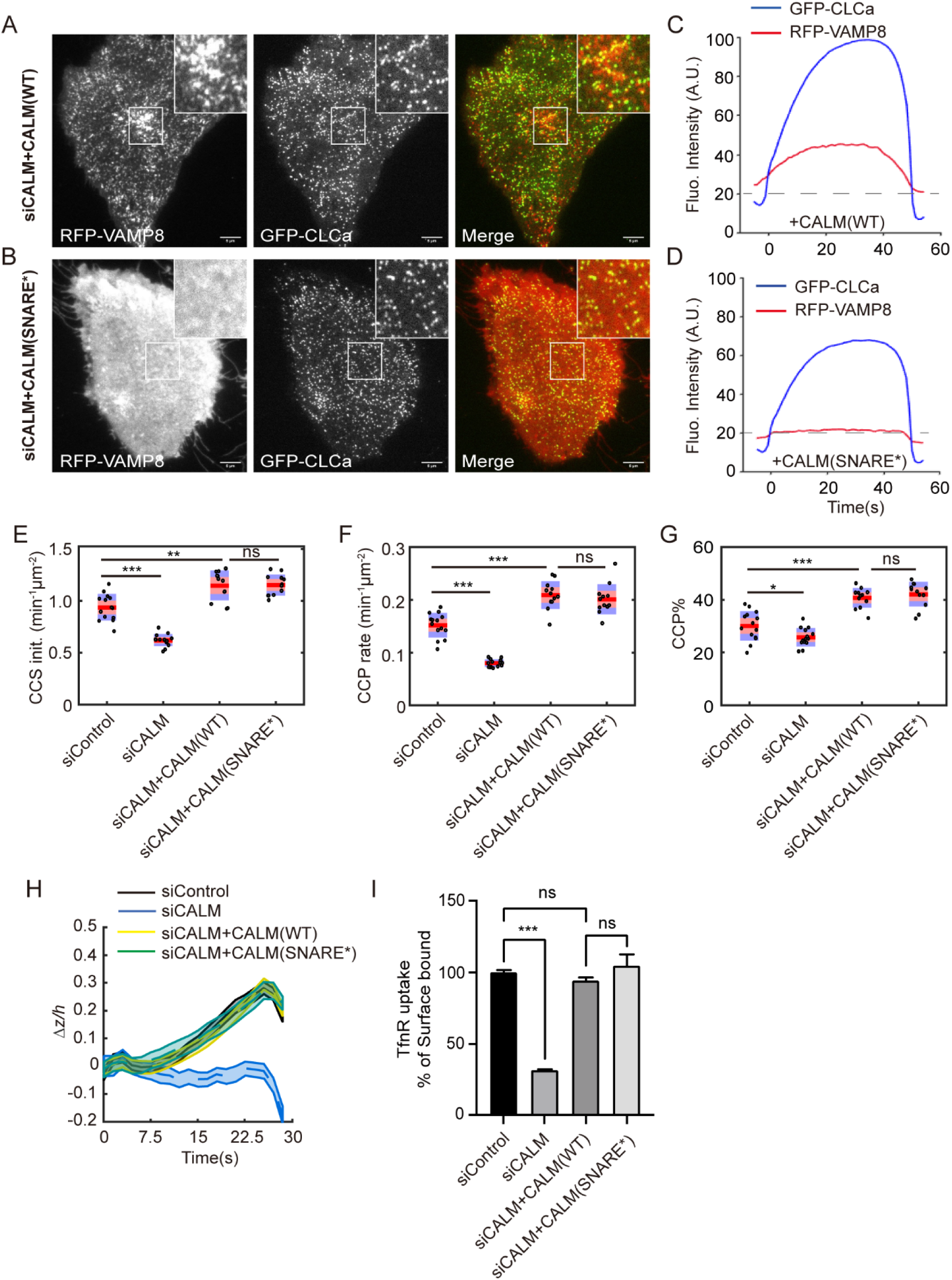
Abolishing VAMP8 recruitment to CCVs does not affect CME. **(A and B)** Representative TIRFM images of ARPE-HPV eGFP-CLCa + RFP-VAMP8 cells that stably express myc-tagged (A) CALM(WT) or (B) CALM(SNARE*) and treated with CALM siRNA to specifically knock down endogenous CALM but not exogenous CALM. White ROI is magnified on the top-right corner. VAMP8 formed clusters on the PM of cells expressing CALM(WT) and was abundant but diffusive on the PM of cells expressing CALM(SNARE*). Scale bars = 5 μm. **(C and D)** The cohort-averaged fluorescence intensity traces for CCPs (tagged with eGFP-CLCa) and CCP-enriched RFP-VAMP8 in cells expressing (C) CALM(WT) or (D) CALM(SNARE*). CALM(SNARE*) failed to recruit VAMP8 to CCPs. Number of dynamic tracks analyzed: 27,028 for (C) and 25,052 for (D). **(E-G)** TIRF microscopy analysis showed that, unlike CALM knockdown, abolishing VAMP8 recruitment to CCVs by rescue with CALM(SNARE*) does not affect (E) CCSs initiation rate, (F) CCPs initiation rate, and (G) % of CCPs. The data presented was acquired from a single experiment that represents three independent repeats. Each dot in (E-G) represents a movie. N = 12 movies for each condition. Number of dynamic tracks analyzed: 200,143 for siControl, 112,579 for siCALM, 140,451 for siCALM+CALM(WT) and 113,132 for siCALM+CALM(SNARE*). **(H)** Epi-TIRF microscopy analysis showed that abolishing VAMP8 recruitment to CCVs does not affect CCP invagination. The data provided were acquired from a single experiment that represents three independent repeats. N = 15 movies for each condition. Shadowed area indicates 95% confidence interval. Number of CCP tracks analyzed to obtain the Δ*z(t)/h* curves: 14,562 for siControl, 2,482 for siCALM, 22,402 for siCALM+CALM(WT) and 16,572 for siCALM+CALM(SNARE*). **(I)** Measurements of the TfnR uptake efficiency. Error bars: SEM of N = 24 samples. Statistical analysis of the data in (E-G) is the Wilcoxon Rank Sum test. Statistical analysis of the data in (I) was performed using GraphPad Prism by unpaired *t*-test. \*\*\**P* ≤ 0.001, \**P* ≤ 0.05, \*\**P* ≤ 0.01, ns, *P* > 0.05.

Next, we compared CCP dynamics and CME efficiency in cells expressing CALM(WT) and CALM(SNARE*) after siRNA-mediated knockdown of endogenous CALM (Fig. S3E). As previously reported (8, 25), depletion of CALM in ARPE-HPV eGFP-CLCa cells significantly reduced CCS initiation rate (Fig. 2E), CCP initiation rate (Fig. 2F), CCP% (Fig. 2G), CCP invagination depth (Fig. 2H), and TfnR uptake efficiency (Fig. 2I), and as expected these defects were fully rescued in cells exogenously expressing siRNA-resistant CALM(WT). Surprisingly, the CALM knockdown phenotypes were also fully rescued by CALM(SNARE*) (Fig. 2E-I), indicating that abolishing VAMP8 recruitment to CCVs does not affect CME.

Together, these unexpected observations suggest that direct recruitment of VAMP8 to CCVs is not required for its role in CME. Therefore, we next explored other possible mechanisms to explain the requirement for VAMP8 in CME.

### VAMP8 knockdown downregulates CME cargo resulting in CME inhibition

Cargo loading has been proposed to be another key factor that determines the fate of CCPs (4, 43–45). We next examined whether VAMP8 knockdown affects the amount of CME cargo.

Differential proteomic analysis of ARPE-HPV cells treated with control or VAMP8 siRNA revealed that VAMP8 knockdown significantly down-regulates several cell surface receptors that are prototypical cargoes of CME (Fig. 3A and Table S1), including TfnR (gene: TFRC), EGFR, VLDLR, and MET. Note that the extent of reduction of surface-expressed receptors is likely attenuated by the inhibition of CME that accompanies VAMP8 knockdown.

**Figure 3.**
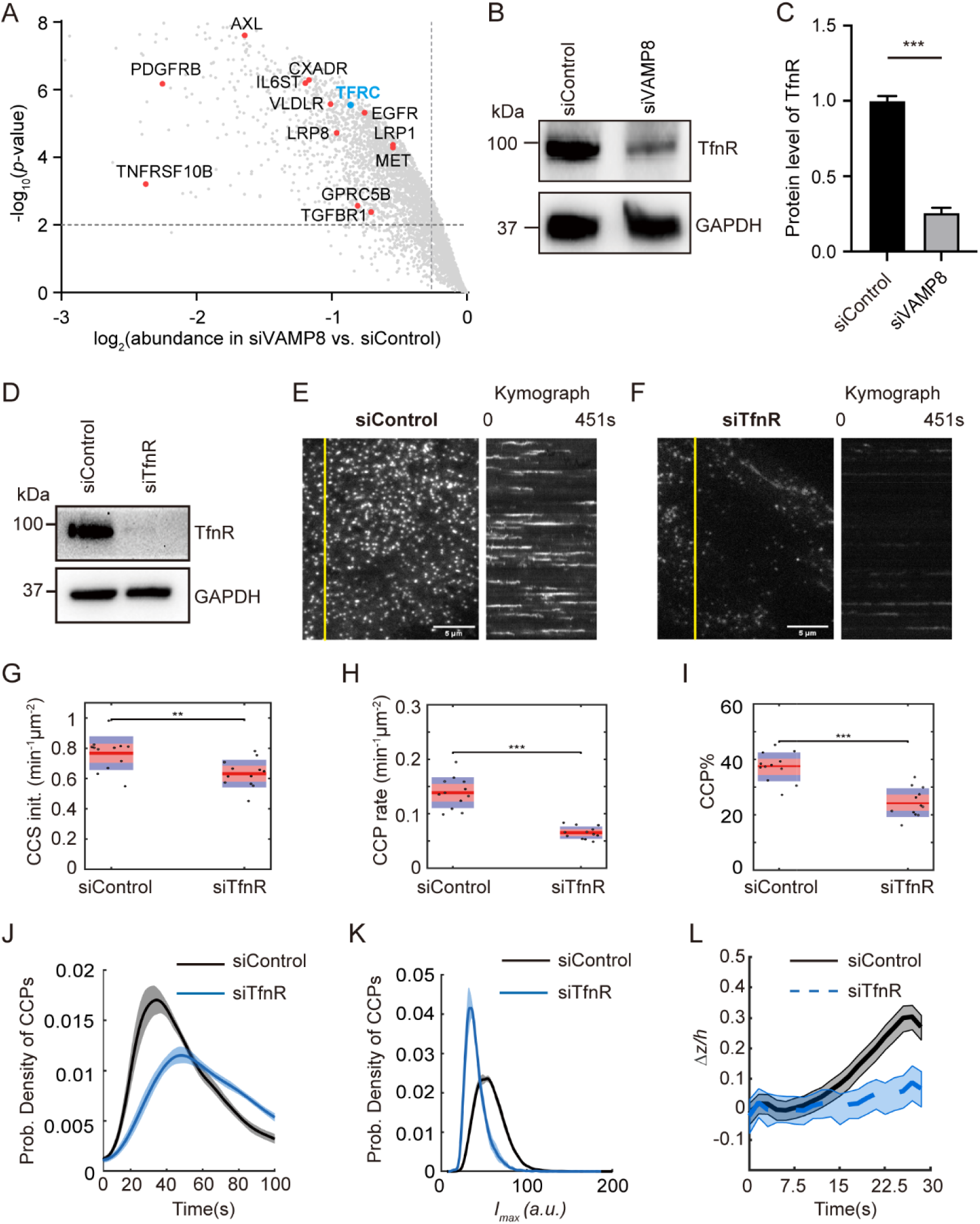
VAMP8 knockdown downregulates CME cargos that results in the inhibition of CME. **(A)** MassSpec analysis revealed that VAMP8 knockdown downregulates the expression levels of several surface receptors (see also in Table S1). Two dashed lines indicate *p*-value = 0.01 and FC = 1.2. **(B)** Western blotting confirmed that VAMP8 knockdown downregulates the level of TfnR in ARPE-HPV eGFP-CLCa cells. **(C)** Quantification of the protein levels of TfnR from the western blotting results. Error bars: SEM of N = 3 biological repeats. **(D)** siRNA treatment efficiently depleted TfnR in ARPE-HPV eGFP-CLCa cells. **(E and F)** Representative single frame images and corresponding kymographs (for region indicated by yellow lines) from TIRFM movies (1 frame/s, 7.5 min/movie) that were captured in ARPE-HPV eGFP-CLCa cells treated with (E) control siRNA or (F) TfnR siRNA (see Movie 3). Scale bars = 5 µm. **(G-I)** Effect of TfnR knockdown on the initiation rates of (G) all CCSs and (H) CCPs, as well as (I) % of CCPs. Each dot represents a movie. **(J and K)** Effect of TfnR knockdown on the (J) lifetime distribution and (K) maximum fluorescence intensity (*I_max_*) distribution of CCPs. The data presented was acquired from a single experiment that represents three independent repeats. N = 12 movies for each condition. Number of dynamic tracks analyzed: 58,038 for siControl and 45,988 for siTfnR. Shadowed area indicates 95% confidence interval. **(L)** Effect of TfnR knockdown on CCP invagination. The data presented was acquired from a single experiment that represents three independent repeats. N = 15 movies for each condition. Number of CCP tracks analyzed to obtain the Δ*z(t)/h* curves: 7,777 for siControl and 741 for siTfnR. 95% confidence interval was indicated by shadow area. Statistical analysis of the data in (C) was performed using GraphPad Prism by unpaired t-test. Statistical analysis of the data in (G-I) is the Wilcoxon Rank Sum test. \*\**P* ≤ 0.01, \*\*\**P* ≤ 0.001.

To determine whether VAMP8 knockdown inhibits CME by reducing the amount of cargo, we selected TfnR for validation and further functional study because: 1) it is an abundant cell surface protein that is primarily internalized via CME; 2) its clustering was observed to promote CCP initiation and increase TfnR uptake (46); and 3) its overexpression led to an increased ratio of CCPs/ACs without affecting their lifetimes, suggesting that its concentration may contribute to the stabilization and maturation of CCPs (45). Western blotting results confirmed that TfnR was downregulated upon VAMP8 knockdown (Fig. 3B and C).

Next, we directly knocked down TfnR in ARPE-HPV eGFP-CLCa cells (Fig. 3D) and strikingly observed that TfnR depletion phenocopies VAMP8 knockdown leading to: i) a strong reduction in the amount of clathrin on the PM (Fig. 3E and F and Movie 3); ii) a significant reduction in CCS initiation rate (Fig. 3G), CCP initiation rate (Fig. 3H) and CCP% (Fig. 3I); iii) extension of CCP lifetimes (Fig. 3J), iv) reduction in the size of CCPs (Fig. 3K); and iv) strong inhibition of CCP invagination depth (Fig. 3L).

Together these results demonstrate that VAMP8 knockdown down-regulates the abundant and prototypical CME cargo, e.g. TfnR, which results in the inhibition of CCP initiation, stabilization and maturation. We next aimed to understand how VAMP8 knockdown down-regulates CME cargo.

### VAMP8 knockdown inhibits endocytic recycling and misdirects TfnR towards degradation in lysosomes

To test whether the downregulation of TfnR is a result of transcriptional inhibition, we conducted RT-qPCR and observed that VAMP8 knockdown did not significantly affect the mRNA level of TfnR (Fig. 4A), suggesting that TfnR was degraded upon VAMP8 knockdown.

**Figure 4.**
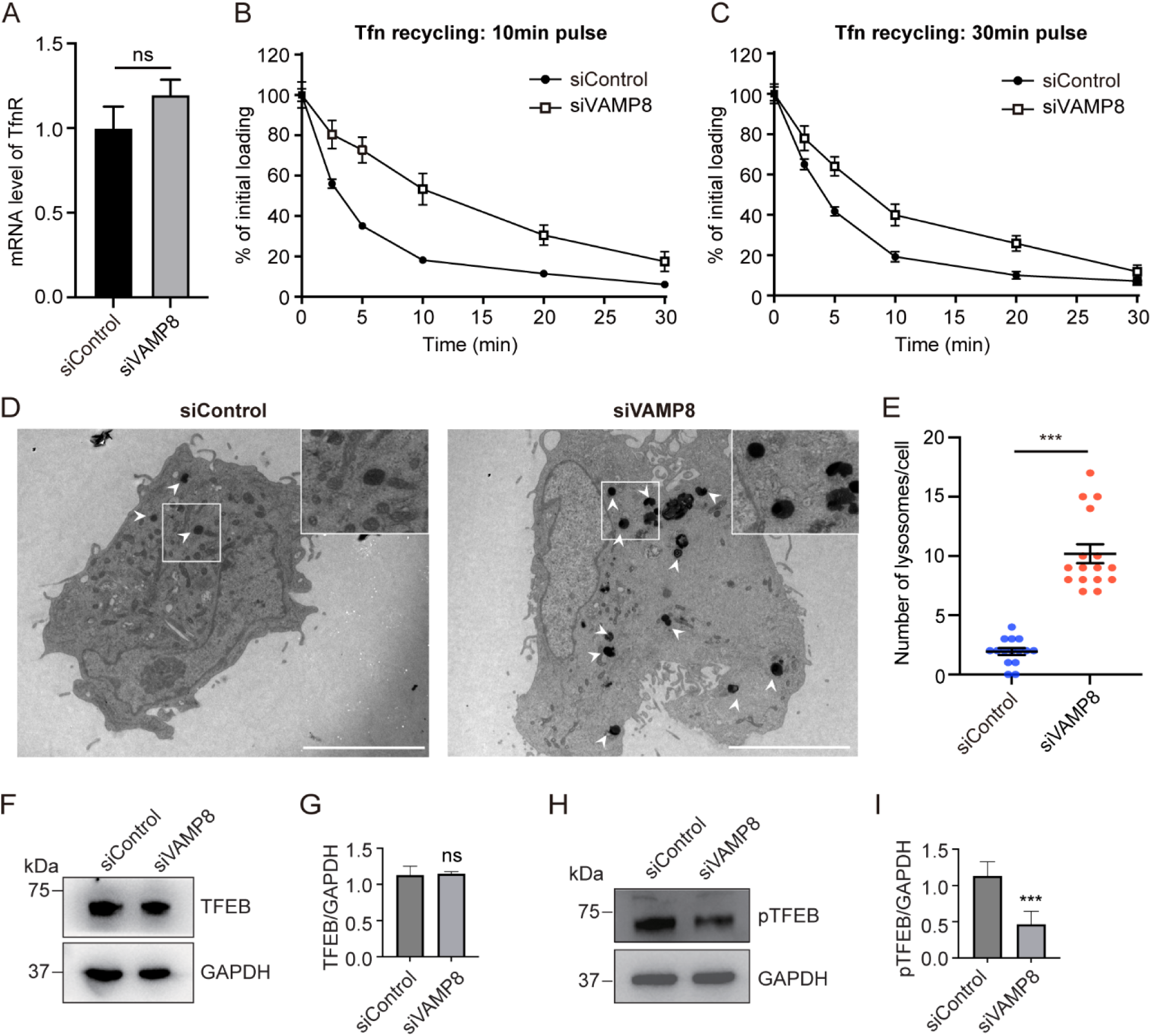
VAMP8 knockdown inhibits endocytic recycling and accumulates lysosomes. **(A)** Quantification of the mRNA level of TfnR. GAPDH mRNA was used as an internal standard for normalization. Error bars: SEM of N = 9 samples. **(B and C)** In-cell ELISA assay revealed that VAMP8 knockdown inhibits Biotin-Tfn recycling. Biotin-Tfn was initially loaded into cells for (B) 10min pulse to measure fast recycling or (C) 30min pulse to measure slow recycling. Error bars: SEM of N = 12 samples. **(D)** Representative transmission electron microscope images of ARPE-HPV eGFP-CLCa cells treated with control (left) or VAMP8 (right) siRNA. Lysosomes accumulated upon VAMP8 knockdown. White ROI is magnified on the top-right corner. White arrowheads point to lysosomes. Scale bars = 5μm. **(E)** Quantification of the number of lysosomes per cell. Each dot represents a cell. Error bars in (E) indicate SEM of N = 15 for siControl and N = 16 for siVAMP8. **(F and G)** Western blotting indicates that VAMP8 knockdown did not affect the protein levels of Transcription factor EB (TFEB). **(H and I)** Western blotting indicates that VAMP8 knockdown reduced TFEB phosphorylation level. Error bars in (G and I) indicate SD of N = 3 biological repeats. Statistical analysis of the data in (A, E, G and I) was performed using GraphPad Prism by unpaired t-test. \*\*\**P* ≤ 0.001, ns, *P* > 0.05.

In normal cells, the bulk of internalized Tfn/TfnR complexes are recycled back to the PM for reutilization (47, 48), while only a very small portion of TfnR is retained in late endosomes and delivered to lysosomes for degradation (49, 50). In vitro studies have demonstrated that VAMP8 mediates homotypic fusion of both early and late endosomes (26, 38, 39), while another study established a role for VAMP8 in the fusion of Rab11-positive recycling endosomes at immunological synapses in activated CD8-positive T cells (34). Although a role for VAMP8 in TfnR recycling has not been reported, based on these finding, we reasonably hypothesized that VAMP8 knockdown could skew endocytic recycling of TfnR towards lysosomal degradation.

To test this hypothesis, we first examined whether the endocytic recycling of Tfn/TfnR was affected by VAMP8 knockdown via measuring the rate of recycling of internalized Tfn following well-established protocols (41, 51). Internalized Tfn/TfnR can be recycled to the PM through two pathways: i) a fast recycling pathway, measured following a brief pulse of Tfn uptake, where Tfn/TfnR is recycled to the PM from early endosomes; and ii) a slow recycling pathway, measured following a prolonged pulse of Tfn uptake, where Tfn/TfnR is first transported to the recycling endosomes before being recycled to the PM (15). Upon siRNA-mediated knockdown of VAMP8, we found that both the fast recycling from the early endosome (Fig. 4B) and the slow recycling from the recycling endosomes (Fig. 4C) of Tfn were significantly inhibited, implying that VAMP8 plays a key role in regulating Tfn/TfnR recycling.

We next examined whether more TfnR was delivered to lysosomes for degradation upon VAMP8 knockdown. Firstly, we conducted electron microscopy imaging and revealed a significant increase in the number of electron-dense lysosomes when VAMP8 was knocked down in ARPE-HPV cells (Fig. 4D and E), which is an indication of enhanced lysosome activities. The increase in lysosome number is consistent with the observed decrease in TFEB phosphorylation level (Fig. 4F-I), indicating enhanced lysosome biogenesis.

Secondly, we treated cells with leupeptin, a well-established inhibitor of lysosomal proteases, and observed by Western blotting that leupeptin treatment rescued the reduction of TfnR levels caused by VAMP8 knockdown (Fig. 5A-C). We next conducted immunofluorescence in cells expressing LAMP1-mCherry, a marker of late endosomes/lysosomes. Consistent with our electron microscopy and biochemical studies, we observed that VAMP8 knockdown increased the overall number of LAMP1-positive endo-lysosomes (Fig. 5D, quantified in E), and decreased the amount of TfnR (Fig. 5D, quantified in F). Moreover, the colocalization of TfnR with LAMP1 was significantly enhanced (Fig. 5D, quantified in G). In control cells treated with leupeptin, the levels of TfnR were unaffected (Fig. 5F); however, leupeptin treatment enhanced the number of LAMP1-positive endo-lysosomes (Fig. 5E) and the degree of colocalization of TfnR (Fig. 5G), presumably a reflection of the normal turnover of TfnR in lysosomes. Leupeptin treatment of VAMP8 knockdown cells successfully restored TfnR levels and significantly enhanced the colocalization of LAMP1 and TfnR (Fig. 5F and G). Interestingly, upon leupeptin treatment in VAMP8 knockdown cells, TfnR and LAMP1 primarily colocalized in tightly clustered lysosomes in the perinuclear regions (Fig. 5D, quantified in H), rendering the depletion of peripherally located TfnR more apparent. As a result, although leupeptin treatment restored the cellular level of TfnR in VAMP8 knockdown cells, the lysosome-trapped, non-degraded TfnR was unable to rescue either the levels of surface TfnR (Fig. S5A) or its uptake efficiency by CME (Fig. S5B).

**Figure 5.**
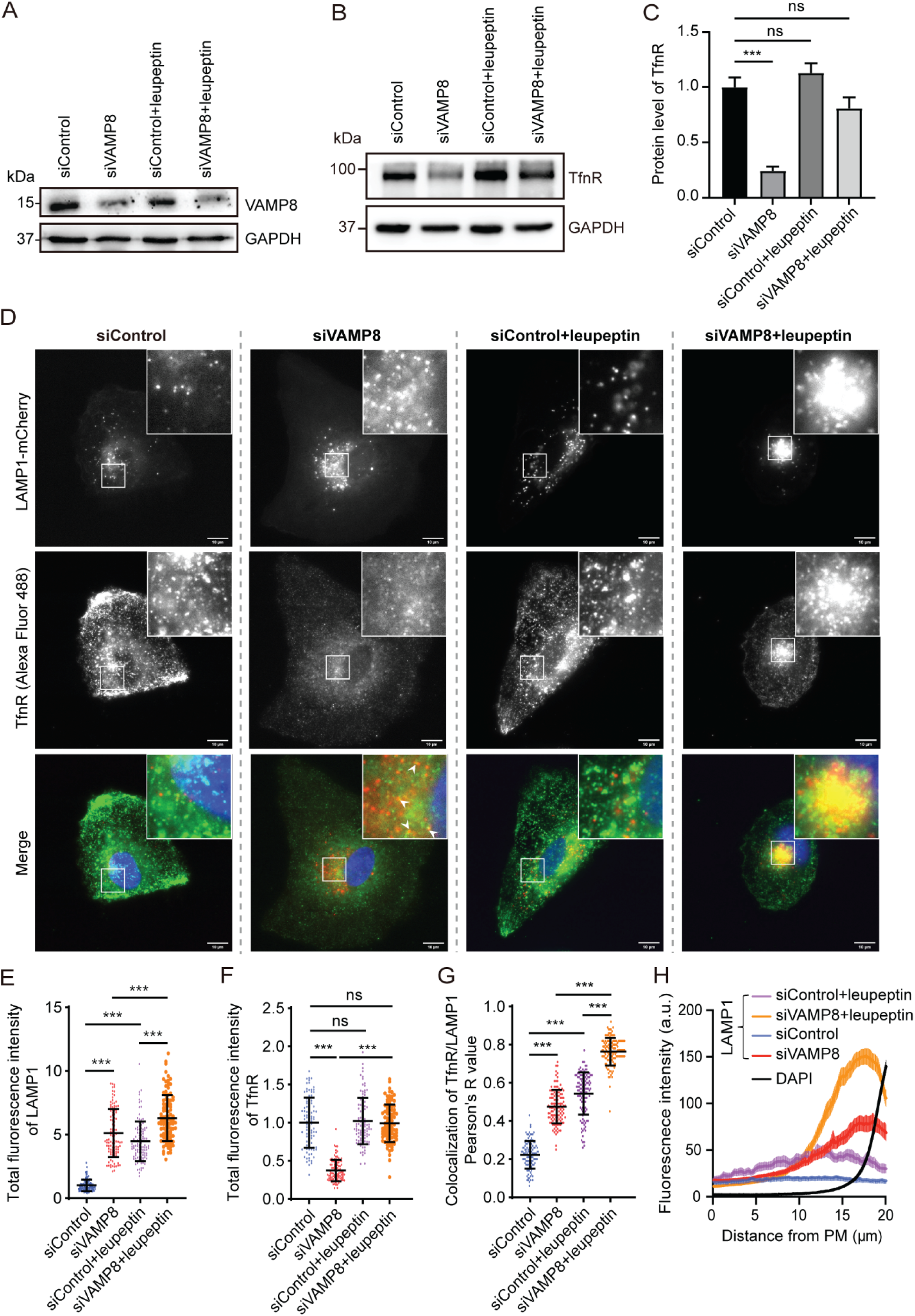
VAMP8 knockdown skews endosomal recycling of TfnR towards lysosomal degradation. **(A-C)** ARPE-HPV eGFP-CLCa cells were treated with 100 μM leupeptin and transfected with indicated siRNAs for 72 h. (A) Western blotting indicates the knockdown efficiency of VAMP8. (B) Western blotting indicates the protein levels of TfnR. (C) Quantification of TfnR levels from western blotting results. Error bars indicate SEM of N = 3 biological repeats. **(D)** Representative Epi-fluorescence images of ARPE-HPV cells that stably express LAMP1-mCherry (red) and were stained with anti-TfnR antibody (green, Alexa Fluor 488-anti-mouse) and DAPI (blue). White ROI is magnified on the top-right corner. White arrowheads point to the colocalized LAMP1 and TfnR. Scale bars = 10 μm. **(E)** The total fluorescence intensities of LAMP1-mCherry. **(F)** The total fluorescence intensities of TfnR. **(G)** Pearson’s correlation coefficients of TfnR and LAMP1. Each dot shown in (E-G) represents a cell. N = 93 for siControl, N = 100 for siVAMP8, N = 101 for siControl+leupeptin, and N = 93 for siVAMP8+leupeptin. Error bars in (E-G) indicate SD. **(H)** Lysosomal perinuclear index was assessed by measuring the fluorescence intensity distributions of LAMP1-mCherry from the PM to nuclear. Intensity traces were obtained by drawing lines from the PM to nucleus using ImageJ. N = 50 cells (2 lines per cell) were counted to generate the mean fluorescence intensity traces. Shadowed area indicates 95% confidential interval. DAPI signals were used as reference. Statistical analysis of the data in (C, E, F and G) was performed using GraphPad Prism by unpaired t-test, *** *P* ≤ 0.001, ns: *P* > 0.05.

Together these observations demonstrate that VAMP8 knockdown impairs Tfn/TfnR recycling from the early and recycling endosomes to the PM, resulting in the mis-sorting of TfnR to late endosomes/lysosomes and its degradation therein (Fig. 6). We conclude that the inhibitory effects of VAMP8 knockdown on early and late stages of CME are indirect and a result of VAMP8’s role in recycling of endocytic cargo and the depletion of cargo molecules from the cell surface (Fig. 6).

**Figure 6.**
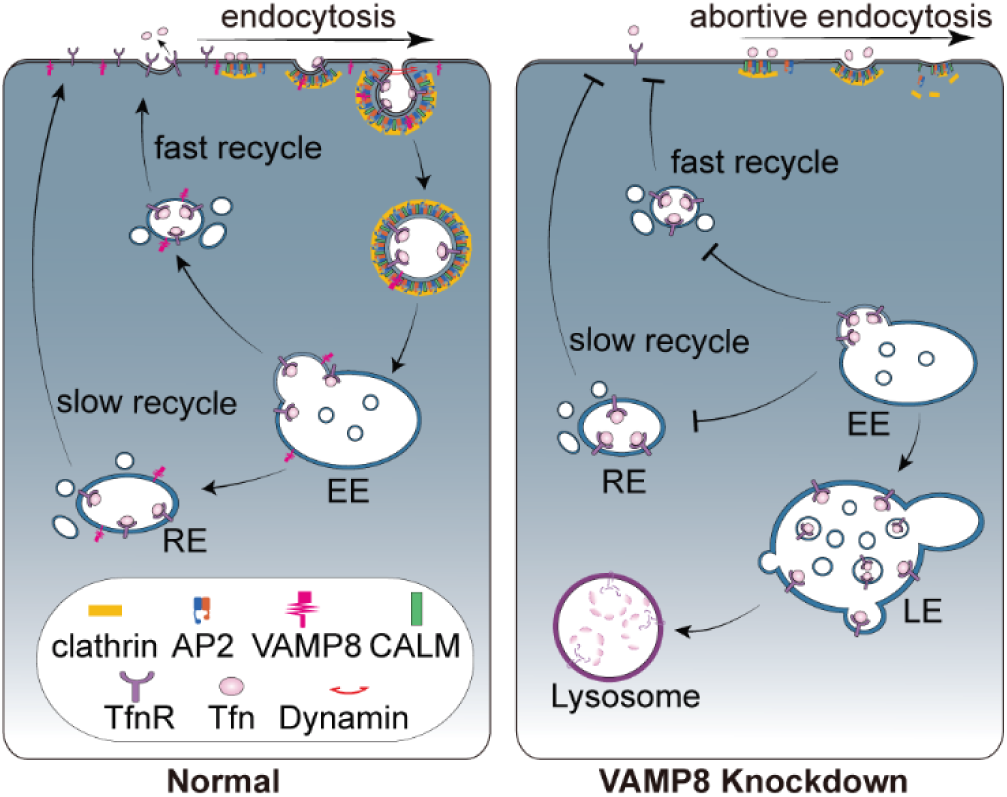
Illustration of intracellular trafficking of Tfn/TfnR complex in the presence (left) and in the absence of VAMP8 (right). EE: early endosome; RE: recycling endosome; LE: late endosome. In normal cells, internalized Tfn/TfnR can be recycled to the PM to maintain CME. Nonetheless, VAMP8 knockdown reroutes Tfn/TfnR to the lysosome for degradation, resulting in the inhibition of CME.

## Discussion

VAMP8 is a ubiquitously expressed R-SNARE protein best studied for its role in regulated exocytosis, exemplified by the defective secretion phenotypes observed in VAMP8 knockout mice (30–33). VAMP8 is also involved in lysosome-autophagosome fusion (29, 35, 52) and the homotypic fusion of early and late endosomes (26, 38, 39). Few studies, however, have explored the role of VAMP8 in CME. Here we show that VAMP8 depletion inhibits CME indirectly through defects in endocytic recycling, diverting endocytic cargo towards lysosomal degradation and leading to a reduction of transmembrane cargo essential for CCP assembly, stabilization, curvature generation and productive CCP formation. These studies reveal that VAMP8 mediates a tight coupling between endocytic recycling and endocytosis, which impacts every stage of CME.

### Re-evaluating the role of SNARE proteins in CME

Previous studies proposed that SNARE incorporation into nascent CCVs might be a prerequisite for CME progression, given their essential role in priming vesicles for downstream fusion events (25). This hypothesis is consistent with evidence for direct roles for SNARE proteins in stabilizing nascent COPI (16) and COPII (17, 18) vesicles. However, we show that abolishing VAMP8 recruitment to CCPs–via expression of the CALM (SNARE*) mutant–does not impair CME (Fig. 2). This result appears to challenge the generalization that SNARE loading directly regulates vesicle formation.

We focused our studies on VAMP8 given its localization to CCPs, that it binds to CALM with the highest affinity relative to other R-SNAREs (20), and its established role in the earliest endosomal fusion events (26). However, it is possible that other R-SNAREs either naturally or in compensation for the loss of CCP-associated VAMP8, might suffice to stabilize CCPs in VAMP8’s absence. Other post-Golgi R-SNAREs include VAMP2, VAMP3, VAMP4 and VAMP7. Notably, VAMP2 and 3 bind, albeit with lower affinity, to the same site on CALM as VAMP8, so that the CALM(SNARE*) would also prevent their recruitment to CCPs. VAMP4 bears a dileucine motif, shown to interact with AP1 (53), but can also be recognized by AP2 (54) and CALM (19); however, unlike VAMP3 and VAMP8, VAMP4 is primarily localized intracellularly (55). VAMP7 is recognized by the clathrin adaptor Hrb (22, 23); however, unlike VAMP3, knockdown of VAMP7 did not phenocopy knockdown of VAMP8 (25). Thus, while we cannot rule out the possibility that the incorporation of SNARE proteins is essential for CCP formation, current data do not support this. Given the multiple functions of SNARE proteins and their adaptors, as well as potential functional redundancy between SNAREs, resolving this question would require simultaneous knockdown of multiple SNAREs and reconstitution studies with discriminating point mutations, as illustrated by (19). It will also be critical to rule out, as we have done here, and indirect effect related to cargo availability.

### VAMP8 links endocytic recycling to CME efficiency

Our data establish that VAMP8 knockdown down-regulates surface TfnR and other CME cargo receptors by disrupting their recycling from early/recycling endosomes to the PM. While VAMP8 is not known to play a role in constitutive exocytosis, it is also possible that a defect in biosynthetic trafficking contributes to cargo depletion. That cargo is rerouted to and degraded in lysosomes, however, suggests that the defect in recycling predominates. Interestingly, the extra burden of normally recycled endocytic receptors imposed on lysosomal degradation resulted in partial activation of the TFEB transcription factor, increased lysosomal biogenesis and increased numbers of lysosomes. Further studies will be necessary to define the mechanism(s) and site(s) at which VAMP8 functions in receptor recycling.

The consequent reduction in cargo availability inhibits CCP dynamics through two primary mechanisms. First, initiation/stabilization of nascent CCPs is impaired. These results are consistent with previous studies showing that cargo, especially cargo with tyrosine-based internalization motifs such as TfnR, play a critical role in determining CME efficiency (43). Indeed, overexpression or clustering of TfnR increases the rate of CCP initiation and the efficiency of CCP maturation (45, 46), and others have shown that CCPs containing a variety of cargo molecules (TfnR, LDLR and/or virus particles) are stabilized relative to vacant CCPs (4). In line with this, our finding that siRNA-mediated depletion of TfnR recapitulated the VAMP8 knockdown phenotype (Fig. 3) further underscores the functional importance of TfnR in this process. Second, and unexpectedly, the ability of CCPs to invaginate is severely impaired when cargo loading is insufficient. The defect in membrane invagination could be due to three, nonexclusive factors: 1) Direct effects of cargo depletion, given that cargo enrichment has been suggested to induce a crowding effect that provides energy for membrane curvature (56–58); 2) Indirect effects of cargo depletion, due to a decrease in cargo-dependent recruitment of peripheral membrane proteins such as AP2 complexes and their associated membrane curvature-generating endocytic accessory factors that contribute to the defect in invagination; 3) Reduced recruitment of CALM, which is known to play a role in curvature generation at CCPs, given the significant contribution of SNARE-CALM interactions in recruiting CALM to the plasma membrane (24).

Importantly, our findings provide direct evidence for the essential role of cargo in CCP formation and the essential role of endocytic recycling in maintaining CME. Our work positions VAMP8-dependent cargo recycling as a critical checkpoint for CME. By controlling cargo fate, VAMP8 integrates endocytic uptake with post-endocytic sorting, ensuring a steady supply of receptors to the PM and sustaining CME in non-neuronal cells, with potential implications for tissue homeostasis, nutrient uptake, and receptor signaling.

## Materials and Methods

### Plasmids

The VAMP8 (full-length, human) cDNA was purchased from Addgene (cat# 92424) and mutated to be siRNA resistant while retaining the amino acid sequence. RFP and VAMP8 were then subcloned into pLVX-IRES-puro vector employing NEBuilder® HiFi DNA Assembly Master Mix (Vazyme, Cat# E2621) to generate the RFP-VAMP8 construct. The LAMP1-mCherry (full-length, human) plasmid was purchased from Addgene (cat# 45147). eGFP-CLCa in pLVX-IRES-puro vector was generated in our previous study (41). The siRNA resistant and myc-tagged CALM (full-length, human) cDNA was a kind gift from David J. Owen (Cambridge, UK) in pBMN vector and then mutated (SNARE*: L219S/M244K) to abolish CALM-VAMP8 interactions. Primers used for cloning and mutagenesis are listed in Table S2.

### Antibodies

Primary antibodies used in this study include rabbit polyclonal antibody to VAMP8 (Cell Signaling Technology, 13060S), rabbit polyclonal antibody to CALM (Proteintech, 28554-1-AP), rabbit polyclonal antibody to TfnR (AiFang biological, AF301044) and rabbit polyclonal antibody to GAPDH (Abclonal, AC1001). Secondary antibody used in Electrochemiluminescence (ECL) is HRP-conjugated Goat-anti-rabbit (Abclonal, AS014). Secondary antibodies used in the immunofluorescence assay include Alexa Fluor 488-anti-mouse (Jackson, 170970), and Alexa Fluor 647-anti-mouse (Cell Signaling Technology, 4410S).

### Cell culture and siRNA transfection

HEK 293T and ARPE19-HPV16 (referred to as ARPE-HPV) cells were purchased from ATCC and cultured in DMEM (Gibco, Cat# 8122070) and DMEM/F12 (Gibco, Cat# 8122502), respectively. Culture medium was supplemented with 10% FBS.

For siRNA transfection, cells were seeded on 6-well plates overnight to reach ∼50% confluence before siRNA treatment. Two rounds of siRNA treatment were performed through three days. For each round of siRNA treatment, 0.11 nmol siRNA was added and the siRNA transfection was mediated by Opti-MEM and Lipofectamine RNAi-MAX (Invitrogen, cat# 13778150). Negative Control siRNA was purchased from Thermo Fisher (Silencer Select Negative Control #1 siRNA, cat# 4390843). The siRNAs targeting VAMP2, VAMP8 and transferrin receptor (TfnR) were reported in previous studies and synthesized by GenePharma. siVAMP2: 5’-CAAGCGCAGCCA AGCUC-3’; siVAMP8 #1: 5’-CGCAACAAGACAGAGGATCTGGAAG-3’; siVAMP8 #2: mixture of 5’-AGGAAAUGAUCGUGUGCGGAACCU-3’ and 5’-GGCUCGAAAAUUCUGGU GGAAGAA-3’; siCALM: 5’-ACAGUUGGCAGACAGUUU A-3’; siTfnR: 5’-AACUUCAAGG UUUCUGCCAGC-3’ (20, 59, 60). For cells treated with leupeptin, 100 µM leupeptin (MCE, cat# HY-18234A) was added at the beginning of the first round of siRNA treatment and maintained for three days.

### Virus production and transfection

Lentiviruses were generated following standard protocol (61). Briefly, HEK293T packaging cells were seeded on 10 cm plate and incubated overnight to reach approximately 60-70% confluency. Next day, the culture medium was replaced with 7 ml fresh culture medium supplemented with pre-mixed NaCl (120 µL from 150 mM filter sterilized stock), RFP-VAMP8 (WT) or LAMP1-mCherry plasmid (5 µg), pSPAX2 packaging vector (3 µg), pMD2G packaging vector (3 µg), and PEI (33 µL from 1 mg/ml stock). Lentiviruses encoding RFP-VAMP8 (WT) or LAMP1-mCherry were harvested after 48-72h and subsequently used to transfect ARPE-HPV eGFP-CLCa cells and ARPE-HPV cells with the assistance of polybrene (Merck, cat# TR-1003-G), respectively. The resulting ARPE-HPV eGFP-CLCa RFP-VAMP8 cells and ARPE-HPV LAMP1-mCherry cells were sorted by Fluorescence-activated Cell Sorting (FACS).

Retroviruses encoding CALM (WT) and CALM (SNARE*) were generated following standard protocol (62). The procedure was the same as described for lentivirus production except that retroviruses were packaged with GAG/POL and VSVG envelopes. The harvested retroviruses were used to transfect ARPE-HPV eGFP-CLCa RFP-VAMP8 cells, ultimately generating ARPE-HPV eGFP-CLCa RFP-VAMP8 CALM(WT) or ARPE-HPV eGFP-CLCa RFP-VAMP8 CALM(SNARE*) cells.

### Transferrin receptor (TfnR) uptake assay

Internalization of TfnR was assessed using in-cell enzyme-linked immunosorbent assay (ELISA) following established protocols (41, 63, 64). Briefly, cells were seeded on 96-well Strip plate (Costar, cat# 11621042) overnight to reach 60%-70% confluency. Before assay, cells were starved for 30 min in 37 °C incubator with pre-warmed PBS^4+^ (1× PBS buffer plus 1 mM CaCl_2_, 1 mM MgCl_2_, 0.2% BSA, and 5 mM D-glucose) and then moved to 4 °C fridge before adding ice-cold PBS^4+^ that contained 5 ug/ml HTR-D65 (a polyclonal antibody to TfnR) (65). Next, cells were divided into three groups: Surface-bound, Blank, and 10 min uptake. Surface bound group cells were kept in 4 °C fridge to measure the amount of surface-bound TfnR. Blank group cells were kept in 4 °C fridge for 10min and then acid-washed (0.2 M acetic acid and 0.2 M NaCl, pH 2.3, 4 °C) to remove surface-bound HTR-D65 that indicates the background signal. 10 min uptake group cells were incubated at 37 °C water bath for 10 min to measure the internalization of TfnR and then acid-washed to remove surface-bound HTR-D65. Subsequently, all groups were washed with cold PBS and fixed with 4% PFA before being permeabilized with 0.1% Triton X-100. Permeabilized cells were blocked with Q-PBS (1×PBS plus 2% BSA, 0.1% lysine, and 0.01% saponin, pH 7.4) and then probed with HRP-conjugated Goat anti-Mouse IgG (BioRad). Finally, O-phenylenediamine dihydrochloride (OPD, Sigma-Aldrich) was added to detect the amount of surface and internalized HTR-D65 while BCA assays were used to account for well-to-well cell number variations.

### Transferrin (Tfn) recycle assay

Recycling of Tfn was assessed following established protocol (41, 51). Before assays, cells were cultured in biotin-free medium for at least 48 h and then seeded on 96-well strip plate overnight to reach a confluency of 60-70%. Next day, cells were starved in PBS^4+^ for 30 min at 37 °C and then cooled down to 4 °C before adding ice-cold PBS^4+^ that contained 5 ug/ml Biotin-Tfn (Sigma-Aldrich). Cells were then incubated at 37 °C water bath for a 10 or 30 min pulse to load Biotin-Tfn into cells. Excess Biotin-Tfn were removed by washing with ice-cold PBS^4+^. Some cells were kept at 4 °C as loading control whose surface-bound Biotin-Tfn was masked with avidin and biocytin. The other cells were incubated at 37 °C for indicated time points to measure the remaining amount of Biotin-Tfn in cells. Subsequently, all cells were acid-washed, fixed, permeabilized, and blocked as stated in TfnR uptake assay. Finally, the amount of Biotin-Tfn was probed with streptavidin-HRP (Thermo Fisher) and O-phenylenediamine dihydrochloride (OPD, Sigma-Aldrich).

### TIRF and Epi-TIRF microscopy

Prior to imaging, cells were seeded on gelatin-coated coverslips (Merck, cat# 2850-22) or glass-bottom dishes (Ibidi, cat# 81218-800) overnight. For live cell imaging, a Nikon Eclipse Ti2 inverted microscope was equipped with: i) a 100x 1.49 Oil immersion lens; ii) a Prime Back Illuminated sCMOS Camera (Prime BSI, 6.5 x 6.5µm pixel size and 95% peak quantum efficiency); iii) a H-TIRF module for TIRF imaging with a penetration depth fixed at 80 nm; iv) a M-TIRF module for Epifluorescence (Epi) imaging; and v) an Oko-lab Cage Incubator to maintain 37 °C and 5% CO2.

Time-lapse movies (451 frames) were acquired at a rate of 1 frame/s for single-channel TIRF imaging, a rate of 0.5 frame/s for dual-channel TIRF imaging, and a rate of 0.67 frame/s for Epi-TIRF imaging. cmeAnalysis (9, 42) and DASC were applied to analyze the imaging data. Detailed information is available on the website: https://github.com/DanuserLab/cmeAnalysis.

### Immunofluorescence

Cells were seeded on gelatin-coated coverslips overnight to reach a confluency of ∼70% and then washed with PBS before fixing with 4% PFA for 30min at room temperature. The fixed cells were permeabilized with 0.1% Triton X-100 for 10 min and then blocked with culture medium for 2 h at room temperature. After immunostaining with primary and secondary antibodies. The cells were imaged with Nikon Eclipse Ti2 inverted microscope and Zeiss LSM980 confocal laser scanning microscope.

### Western Blotting

Cells were harvested and lysed in lysis buffer. Lysates were boiled for 10 min, separated by SDS–PAGE, and transferred to PVDF membranes. Membranes were blocked with 5% non-fat milk in TBST, followed by incubation with primary antibodies at 4 °C overnight and HRP-conjugated secondary antibodies at room temperature for 1 h. Immunoreactive bands were detected using enhanced chemiluminescence (ECL). Quantitative analysis was performed using ImageJ software. The intensity of each target protein band was normalized to the corresponding loading control. Data are representative of at least three independent experiments and are presented as relative values compared with the control group.

### RNA extraction and quantitative reverse transcription polymerase chain reaction (RT-qPCR) assay

Cells were lysed using TransZol UP (Transgen) at room temperature for 5 min and then moved to 1.5 ml centrifuge tube, followed by adding chloroform and shaking vigorously for 30 s before centrifuging at 10,000 × g for 15 min. Isopropanol was added to the supernatant and gelatinous precipitate was obtained after centrifugation at 10,000 × g for 10 min. Next, 75% alcohol was used to wash the gelatinous precipitate by violently shaking. RNA was collected after centrifugation at 7,500 × g for 5 min finally dissolved in RNase-free water. 2 ug RNA was reverse transcribed to the first chain of cDNA in 20 µL reaction system using FastKing-RT Super Mix kit (TIANGEN, cat# Y1629) according to manufacturer’s instructions. The primers used in qPCR were: 5’-CAGGAGGCATTGCTGATGAT-3’ and 5’-GAAGGCTGGGGCTCATTT-3’ for GAPDH; 5’-AGGTCAAAGACAGCGCTCAA-3’ and 5’-GCCACATAACCCCCAGGATT-3’ for TfnR; 5’-TGTGCGGAACCTGCAAAGT −3’ and 5’-CTTCTGCGATGTCGTCTTGAA-3’ for VAMP8.

### Electron microscopy

Cell precipitation was fixed overnight in 2.5% glutaraldehyde and 2% PFA in 0.1 M phosphate buffer (pH 7.2–7.4), then replaced by 1% osmium tetroxide (Ted Pella, Cat# 20816-12-0) at 0°C for 1h. After fixation, samples were stained with 1% aqueous uranyl acetate (Electron Microscopy Sciences, Cat# 22400, USA) for 1 h, then dehydrated in ethanol (Sinopharm, Cat# 64-17-5) and embedded by EMbed 812 (Electron Microscopy Sciences, Cat# 14900, 13710, 19000 and 13600). Ultrathin sections were stained with Sato’s lead citrate and imagined with a HT7800 electron microscope (Hitachi Hi-tech, Hitachinaka-shi, Japan).

### Proteomic Analysis

#### Sample preparation

Cells were seeded on 15 cm plates overnight to reach ∼50% confluency before siRNA treatment. Two rounds of siRNA transfection were performed over three days. After siRNA treatment, the cells lysates were collected and lysed in ten volumes of modified SDT buffer (0.1 M Tris-HCL, pH 7.6, 0.1 M DTT, 1% SDS, 1% SDC) and incubated at 95 °C for 5 min. The lysate was sonicated to shear genomic DNA, and clarified by centrifugation at 20,000 × g for 15 min at 20 °C. The supernatant was transferred to ultrafiltration units (Millipore, Amicon Ultra 4 Ultracel 10 KD) and centrifuged at 4000 × g for 40 min. After centrifugation, the concentrates were mixed with 2 mL of 50 mM iodoacetamide in UA solution (8 M urea, 100 mM Tris-HCl pH 8.5) and incubated in darkness at room temperature (RT) for 30 min followed by centrifugation for 30 min. After alkylation, the filter units were washed four times with 4 ml UA buffer and two times 4 mL of 50 mM ammonium bicarbonate by centrifugation at 4,000 × g. Proteins were then digested with Lys-C (1:100, w/w, Wako) for 6 h at 37 °C and trypsin (1:50, w/w, Promega) overnight at 37 °C. The resulting peptide mixture was acidified (pH 2.0) with formic acid, loaded onto SepPak tC18 cartridges (Waters), desalted and eluted with 70% acetonitrile. The eluted peptides were lyophilized and stored at −80 °C before analysis.

#### LC-MS/MS analysis

MS experiments were performed on a nanoscale Vanquish Neo UHPLC system (Thermo Fisher Scientific) connected to an Orbitrap Exploris 480 MS equipped with a nanoelectrospray source (Thermo Fisher Scientific). Mobile phase A contained 0.1% formic acid (v/v) in water; mobile phase B contained 0.1% formic acid in 80% acetonitrile (ACN). The peptides were dissolved in 0.1% formic acid (FA) with 2% acetonitrile and separated on a RP-HPLC analytical column (75 μm × 25 cm) packed with 2 μm C18 beads (Thermo Fisher Scientific) using a linear gradient ranging from 8% to 28% ACN in 70 min and followed by a linear increase to 44% B in 15 min at a flow rate of 300 nL/min. The Orbitrap Exploris 480 MS acquired data in a data independent manner alternating between full scan MS and MS2 scans. The spray voltage was set at 2.2 kV and the temperature of ion transfer capillary was 300 °C. The MS spectra (380–980 m/z) were collected with 120,000 resolutions, AGC of 3 × 106, and 25 ms maximal injection time. Ions were sequentially fragmented by HCD with 30% normalized collision energy, specified isolated windows 6.0 m/z, 30,000 resolutions. and 54 ms maximal injection time were used.

#### Mass spectrometry data analysis

Raw data were processed using DIA-NN (version 2.0), and MS/MS spectra were searched against the SwissProt human database (downloaded in April 2024), and the total number of entries is 20,354. All searches were carried out with precursor mass tolerance of 10 ppm, oxidation (Met) (+15.9949 Da), carbamidomethylation (Cys) (+57.0215 Da), and acetylation (protein N-terminus) (+42.0106 Da) as modifications, two trypsin missed cleavages allowed when searching for proteins. The peptide and protein identifications were filtered by DIA-NN to control the false discovery rate (FDR) < 1%.

### Statistical analysis

Statistical analysis of the parameters from TIRF and Epi-TIRF microscopy, such as the initiation rates of all CCSs and CCPs and the CCP%, is the Wilcoxon Rank Sum test by Matlab. Statistical analysis of the data obtained from western blotting, TfnR uptake assay, immunofluorescence, RT-qPCR and EM are student’s t-test using GraphPad Prism 8. The data from immunofluorescence are presented as means ± SD, while the data from western blotting, TfnR uptake assay, immunofluorescence, RT-qPCR and EM are presented as means ± SEM. Statistical significance levels: ns: *P* > 0.05, * *P* ≤ 0.05, ** *P* ≤ 0.01, *** *P* ≤ 0.001.

## Data availability

The data supporting the findings of this work are available within the paper and the Supplementary Information files.

## Acknowledgments

We thank Dr. Guan-Yu Xiao (U. Kentucky) for helpful discussions. This work is supported by the National Natural Science Foundation of China [Grant No. 32200564 to Z.C.], the Natural Science Foundation of Hunan Province, China [Grant No. 2024JJ2045 to Z.C.] and the Science and Technology Innovation Program of Hunan Province, China [Grant No. 2021SK1014 to D.W.].

## Author contributions

Z.C., S.L.S., A.L., Y.L., X.L., Y.H., X.W., S.L. and D.W. designed the experiments; A.L., Y.L., Z.H., W.C., Y.H. and P.X. performed the experiments; Z.C., A.L., G.H., D.W. and Y.L. analyzed and interpreted the data; Z.C., S.L.S., A.L. and Y.L. wrote the manuscript with contributions from other co-authors; Z.H. drew the cartoon illustrations.

## References

1. H. T. McMahon, E. Boucrot, Molecular mechanism and physiological functions of clathrin-mediated endocytosis. Nat Rev Mol Cell Biol 12, 517–533 (2011).

2. M. Mettlen, P. H. Chen, S. Srinivasan, G. Danuser, S. L. Schmid, Regulation of Clathrin-Mediated Endocytosis. Annu Rev Biochem 87, 871–896 (2018).

3. M. Kaksonen, A. Roux, Mechanisms of clathrin-mediated endocytosis. Nat Rev Mol Cell Biol 19, 313–326 (2018).

4. M. Ehrlich et al., Endocytosis by random initiation and stabilization of clathrin-coated pits. Cell 118, 591–605 (2004).

5. Z. He et al., Dynamic early recruitment of GAK-Hsc70 regulates coated pit maturation. Proc Natl Acad Sci U S A 122, e2503738122 (2025).

6. Z. Yang, et al. (2025) CCDC32 stabilizes clathrin-coated pits and drives their invagination. (eLife Sciences Publications, Ltd).

7. D. Loerke, M. Mettlen, S. L. Schmid, G. Danuser, Measuring the Hierarchy of Molecular Events During Clathrin-Mediated Endocytosis. Traffic 12, 815–825 (2011).

8. X. Wang et al., DASC, a sensitive classifier for measuring discrete early stages in clathrin-mediated endocytosis. Elife 9, e53686 (2020).

9. F. Aguet, Costin N. Antonescu, M. Mettlen, Sandra L. Schmid, G. Danuser, Advances in Analysis of Low Signal-to-Noise Images Link Dynamin and AP2 to the Functions of an Endocytic Checkpoint. Developmental Cell 26, 279–291 (2013).

10. Y. A. Chen, R. H. Scheller, SNARE-mediated membrane fusion. Nat Rev Mol Cell Biol 2, 98–106 (2001).

11. D. Fasshauer, R. B. Sutton, A. T. Brunger, R. Jahn, Conserved structural features of the synaptic fusion complex: SNARE proteins reclassified as Q- and R-SNAREs. Proc Natl Acad Sci U S A 95, 15781–15786 (1998).

12. W. Hong, SNAREs and traffic. Biochim Biophys Acta 1744, 120–144 (2005).

13. I. Dingjan et al., Endosomal and Phagosomal SNAREs. Physiol Rev 98, 1465–1492 (2018).

14. R. Jahn, R. H. Scheller, SNAREs--engines for membrane fusion. Nat Rev Mol Cell Biol 7, 631–643 (2006).

15. F. R. Maxfield, T. E. McGraw, Endocytic recycling. Nat Rev Mol Cell Biol 5, 121–132 (2004).

16. U. Rein, U. Andag, R. Duden, H. D. Schmitt, A. Spang, ARF-GAP-mediated interaction between the ER-Golgi v-SNAREs and the COPI coat. J Cell Biol 157, 395–404 (2002).

17. E. Mossessova, L. C. Bickford, J. Goldberg, SNARE selectivity of the COPII coat. Cell 114, 483–495 (2003).

18. S. Springer, R. Schekman, Nucleation of COPII vesicular coat complex by endoplasmic reticulum to Golgi vesicle SNAREs. Science 281, 698–700 (1998).

19. D. A. Sahlender, P. Kozik, S. E. Miller, A. A. Peden, M. S. Robinson, Uncoupling the functions of CALM in VAMP sorting and clathrin-coated pit formation. PLoS One 8, e64514 (2013).

20. S. E. Miller et al., The molecular basis for the endocytosis of small R-SNAREs by the clathrin adaptor CALM. Cell 147, 1118–1131 (2011).

21. S. J. Koo et al., SNARE motif-mediated sorting of synaptobrevin by the endocytic adaptors clathrin assembly lymphoid myeloid leukemia (CALM) and AP180 at synapses. Proc Natl Acad Sci U S A 108, 13540–13545 (2011).

22. M. Chaineau, L. Danglot, V. Proux-Gillardeaux, T. Galli, Role of HRB in clathrin-dependent endocytosis. J Biol Chem 283, 34365–34373 (2008).

23. P. R. Pryor et al., Molecular basis for the sorting of the SNARE VAMP7 into endocytic clathrin-coated vesicles by the ArfGAP Hrb. Cell 134, 817–827 (2008).

24. Sharon E. Miller et al., CALM Regulates Clathrin-Coated Vesicle Size and Maturation by Directly Sensing and Driving Membrane Curvature. Developmental Cell 33, 163–175 (2015).

25. M. Bhave et al., Functional characterization of 67 endocytic accessory proteins using multiparametric quantitative analysis of CCP dynamics. Proc Natl Acad Sci U S A 117, 31591–31602 (2020).

26. W. Antonin, C. Holroyd, R. Tikkanen, S. Honing, R. Jahn, The R-SNARE endobrevin/VAMP-8 mediates homotypic fusion of early endosomes and late endosomes. Mol Biol Cell 11, 3289–3298 (2000).

27. F. Bilan et al., Endosomal SNARE proteins regulate CFTR activity and trafficking in epithelial cells. Exp Cell Res 314, 2199–2211 (2008).

28. R. D. Bagshaw, D. J. Mahuran, J. W. Callahan, A proteomic analysis of lysosomal integral membrane proteins reveals the diverse composition of the organelle. Mol Cell Proteomics 4, 133–143 (2005).

29. E. Itakura, C. Kishi-Itakura, N. Mizushima, The hairpin-type tail-anchored SNARE syntaxin 17 targets to autophagosomes for fusion with endosomes/lysosomes. Cell 151, 1256–1269 (2012).

30. C. C. Wang et al., A role of VAMP8/endobrevin in regulated exocytosis of pancreatic acinar cells. Dev Cell 7, 359–371 (2004).

31. C. C. Wang et al., VAMP8/endobrevin as a general vesicular SNARE for regulated exocytosis of the exocrine system. Mol Biol Cell 18, 1056–1063 (2007).

32. N. Tiwari et al., VAMP-8 segregates mast cell-preformed mediator exocytosis from cytokine trafficking pathways. Blood 111, 3665–3674 (2008).

33. N. Puri, P. A. Roche, Mast cells possess distinct secretory granule subsets whose exocytosis is regulated by different SNARE isoforms. Proc Natl Acad Sci U S A 105, 2580–2585 (2008).

34. M. R. Marshall et al., VAMP8-dependent fusion of recycling endosomes with the plasma membrane facilitates T lymphocyte cytotoxicity. J Cell Biol 210, 135–151 (2015).

35. J. Diao et al., ATG14 promotes membrane tethering and fusion of autophagosomes to endolysosomes. Nature 520, 563–566 (2015).

36. Q. Chen et al., Prefused lysosomes cluster on autophagosomes regulated by VAMP8. Cell Death & Disease 12, 939 (2021).

37. F. Jian et al., The STX17-SNAP47-VAMP7/VAMP8 complex is the default SNARE complex mediating autophagosome–lysosome fusion. Cell Research 34, 151–168 (2024).

38. W. Antonin et al., A SNARE complex mediating fusion of late endosomes defines conserved properties of SNARE structure and function. EMBO J 19, 6453–6464 (2000).

39. P. R. Pryor et al., Combinatorial SNARE complexes with VAMP7 or VAMP8 define different late endocytic fusion events. EMBO Rep 5, 590–595 (2004).

40. M. Mettlen, G. Danuser, Imaging and modeling the dynamics of clathrin-mediated endocytosis. Cold Spring Harb Perspect Biol 6, a017038 (2014).

41. Z. Chen et al., Wbox2: A clathrin terminal domain-derived peptide inhibitor of clathrin-mediated endocytosis. J Cell Biol 219 (2020).

42. K. Jaqaman et al., Robust single-particle tracking in live-cell time-lapse sequences. Nat Methods 5, 695–702 (2008).

43. Z. Kadlecova et al., Regulation of clathrin-mediated endocytosis by hierarchical allosteric activation of AP2. Journal of Cell Biology 216, 167–179 (2016).

44. H. Maib, F. Ferreira, S. Vassilopoulos, E. Smythe, Cargo regulates clathrin-coated pit invagination via clathrin light chain phosphorylation. J Cell Biol 217, 4253–4266 (2018).

45. D. Loerke et al., Cargo and dynamin regulate clathrin-coated pit maturation. PLoS Biol 7, e57 (2009).

46. A. P. Liu, F. Aguet, G. Danuser, S. L. Schmid, Local clustering of transferrin receptors promotes clathrin-coated pit initiation. J Cell Biol 191, 1381–1393 (2010).

47. M. W. Hentze, M. U. Muckenthaler, N. C. Andrews, Balancing acts: molecular control of mammalian iron metabolism. Cell 117, 285–297 (2004).

48. H. Kawabata, Transferrin and transferrin receptors update. Free Radic Biol Med 133, 46–54 (2019).

49. K. Sirohi et al., M98K-OPTN induces transferrin receptor degradation and RAB12-mediated autophagic death in retinal ganglion cells. Autophagy 9, 510–527 (2013).

50. T. Matsui, T. Itoh, M. Fukuda, Small GTPase Rab12 regulates constitutive degradation of transferrin receptor. Traffic 12, 1432–1443 (2011).

51. P.-H. Chen et al., Crosstalk between CLCb/Dyn1-Mediated Adaptive Clathrin-Mediated Endocytosis and Epidermal Growth Factor Receptor Signaling Increases Metastasis. Developmental Cell 40, 278–288.e275 (2017).

52. S. van Tol et al., VAMP8 Contributes to the TRIM6-Mediated Type I Interferon Antiviral Response during West Nile Virus Infection. J Virol 94 (2020).

53. A. A. Peden, G. Y. Park, R. H. Scheller, The Di-leucine Motif of Vesicle-associated Membrane Protein 4 Is Required for Its Localization and AP-1 Binding*. Journal of Biological Chemistry 276, 49183–49187 (2001).

54. B. T. Kelly et al., A structural explanation for the binding of endocytic dileucine motifs by the AP2 complex. Nature 456, 976–979 (2008).

55. M. Steegmaier, J. Klumperman, D. L. Foletti, J. S. Yoo, R. H. Scheller, Vesicle-associated membrane protein 4 is implicated in trans-Golgi network vesicle trafficking. Mol Biol Cell 10, 1957–1972 (1999).

56. Z. Chen, E. Atefi, T. Baumgart, Membrane Shape Instability Induced by Protein Crowding. Biophys J 111, 1823–1826 (2016).

57. W. T. Snead et al., Membrane fission by protein crowding. Proceedings of the National Academy of Sciences 114, E3258–E3267 (2017).

58. J. C. Stachowiak et al., Membrane bending by protein-protein crowding. Nat Cell Biol 14, 944–949 (2012).

59. H. Huang et al., mTOR-mediated phosphorylation of VAMP8 and SCFD1 regulates autophagosome maturation. Nat Commun 12, 6622 (2021).

60. I. Nakase et al., Transferrin receptor-dependent cytotoxicity of artemisinin-transferrin conjugates on prostate cancer cells and induction of apoptosis. Cancer Lett 274, 290–298 (2009).

61. R. H. Kutner, X. Y. Zhang, J. Reiser, Production, concentration and titration of pseudotyped HIV-1-based lentiviral vectors. Nat Protoc 4, 495–505 (2009).

62. R. Mann, R. C. Mulligan, D. Baltimore, Construction of a retrovirus packaging mutant and its use to produce helper-free defective retrovirus. Cell 33, 153–159 (1983).

63. S. Srinivasan et al., A noncanonical role for dynamin-1 in regulating early stages of clathrin-mediated endocytosis in non-neuronal cells. PLoS Biol 16, e2005377 (2018).

64. S. D. Conner, S. L. Schmid, Differential requirements for AP-2 in clathrin-mediated endocytosis. J Cell Biol 162, 773–779 (2003).

65. S. L. Schmid, E. Smythe, Stage-specific assays for coated pit formation and coated vesicle budding in vitro. J Cell Biol 114, 869–880 (1991).

