## Supplementary information for "VAMP8 function reveals tight linkage between endocytic recycling and endocytosis"

**Movie 1:** Time-lapse TIRFM imaging was performed on ARPE-HPV eGFP-CLCa cells treated with siControl, siVAMP8 #1, and siVAMP8 #2. Images were captured at a rate of 1 frame per second over a period of 7.5 min. The movie is displayed at 25 times the actual speed.

**Movie 2:** Time-lapse TIRFM imaging was performed on ARPE-HPV eGFP-CLCa cells treated with siControl and siVAMP2. Images were captured at a rate of 1 frame per second over a period of 7.5 min. The movie is displayed at 25 times the actual speed.

**Movie 3:** Time-lapse TIRFM imaging was performed on ARPE-HPV eGFP-CLCa cells treated with siControl and siTfnR. Images were captured at a rate of 1 frame per second over a period of 7.5 min. The movie is displayed at 25 times the actual speed.

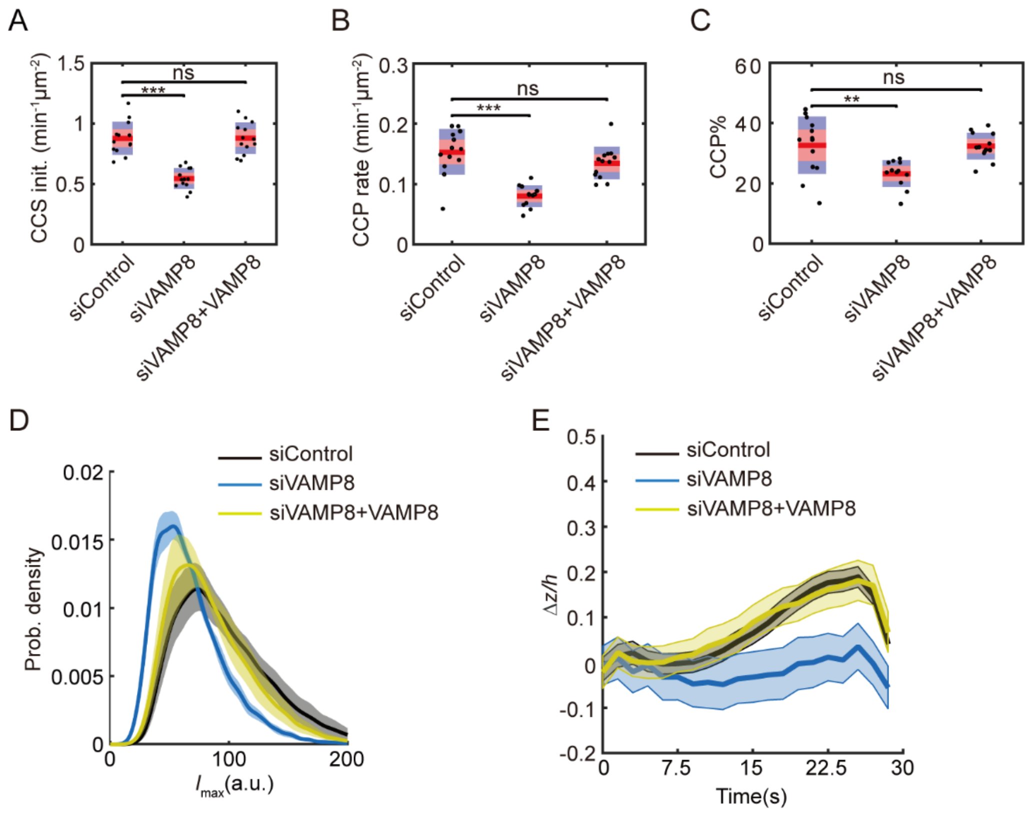

**Figure S1. Expression of siRNA-resistant VAMP8 rescues the phenotypes of VAMP8 knockdown.** Cells were transfected with control or VAMP8 siRNA. **(A-D)** TIRFM analysis revealed that stable expression of siRNA-resistant VAMP8 rescued (A) initiation rate of CCSs, (B) initiation rate of CCPs, (C) % of CCPs, and (D) maximum fluorescence intensities of CCPs. The data presented was obtained from a single experiment (N = 13 movies for each condition). Number of dynamic tracks analyzed: 175,690 for siControl, 120,011 for siVAMP8 and 104,087 for siVAMP8+VAMP8. **(E)** Epi-TIRF microscopy analysis revealed that stable expression of siRNA-resistant VAMP8 rescued the invagination of CCPs. Data presented was obtained from a single experiment (N = 13 movies for siControl, N = 12 movies for siVAMP8, N = 15 movies for siVAMP8+VAMP8). Number of CCP tracks analyzed to obtain the $\Delta z(t)/h$ curves: 20,730 for siControl, 1,567 for siVAMP8 and 4,363 for siVAMP8+VAMP8. Statistical analysis of the data in (A-C) is the Wilcoxon Rank Sum test. ns: $P > 0.05$, $^{**}P \leq 0.01$, $^{***}P \leq 0.001$. Shadowed area in (D-E) indicates 95% confidence interval.

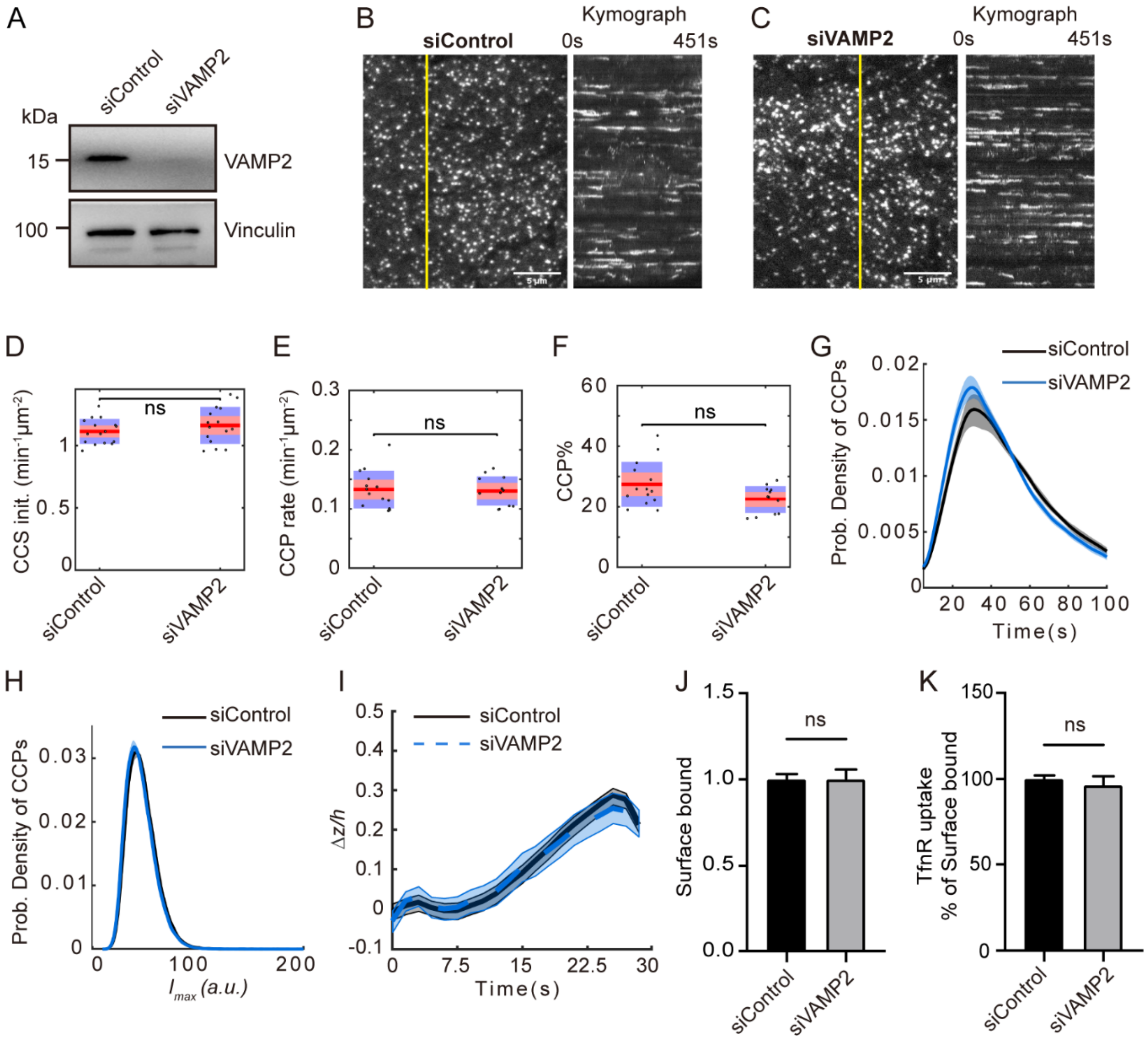

**Figure S2. VAMP2 knockdown does not inhibit CME. (A)** ARPE-HPV eGFP-CLCa cells were transfected with control or VAMP2 siRNA. Western blotting indicates good knockdown efficiency of VAMP2. **(B and C)** Representative single frame images and corresponding kymographs (for region indicated by yellow lines) from TIRFM movies (1 frame/s, 7.5 min/movie, see Movie 2) that were captured in ARPE-HPV eGFP-CLCa cells treated with (B) control or (C) VAMP2 siRNA. Scale bars = 5 µm. **(D-F)** Effect of VAMP2 knockdown on the initiation rates of (D) all CCSs and (E) CCPs, as well as (F) % of CCPs. Each dot represents a movie, and N = 15 movies for each condition. **(G and H)** Effect of VAMP2 knockdown on the (G) lifetime distribution and (H) Maximum fluorescence intensity ($I_{max}$) distribution of CCPs. The data presented was acquired from a single experiment that represents three independent biological repeats. Number of dynamic tracks analyzed: 189,191 for siControl, and 134,710 for siVAMP2. **(I)** Epi-TIRF microscopy analysis showed that VAMP2 knockdown did not affect CCP invagination. The presented data was acquired from a single experiment that represents three independent repeats. N = 15 movies for each condition. Number of CCP tracks analyzed to obtain the $\Delta z(t)/h$ curves: 18,625 for siControl and 9,947 for siVAMP2. Shadowed area in (G-I) indicates 95% confidence interval. **(J and K)** Evaluation of the (J) surface bound and (K) uptake efficiency of TfnR (Internalized/Surface-bound). Error bars in (J and K) indicate SEM of N = 12 samples. Statistical analysis of the data in (J and K) was performed using GraphPad Prism 8 by unpaired t-test: ns, $P > 0.05$. Statistical analysis of the data in (D-F) is the Wilcoxon Rank Sum test, ns: $P > 0.05$.

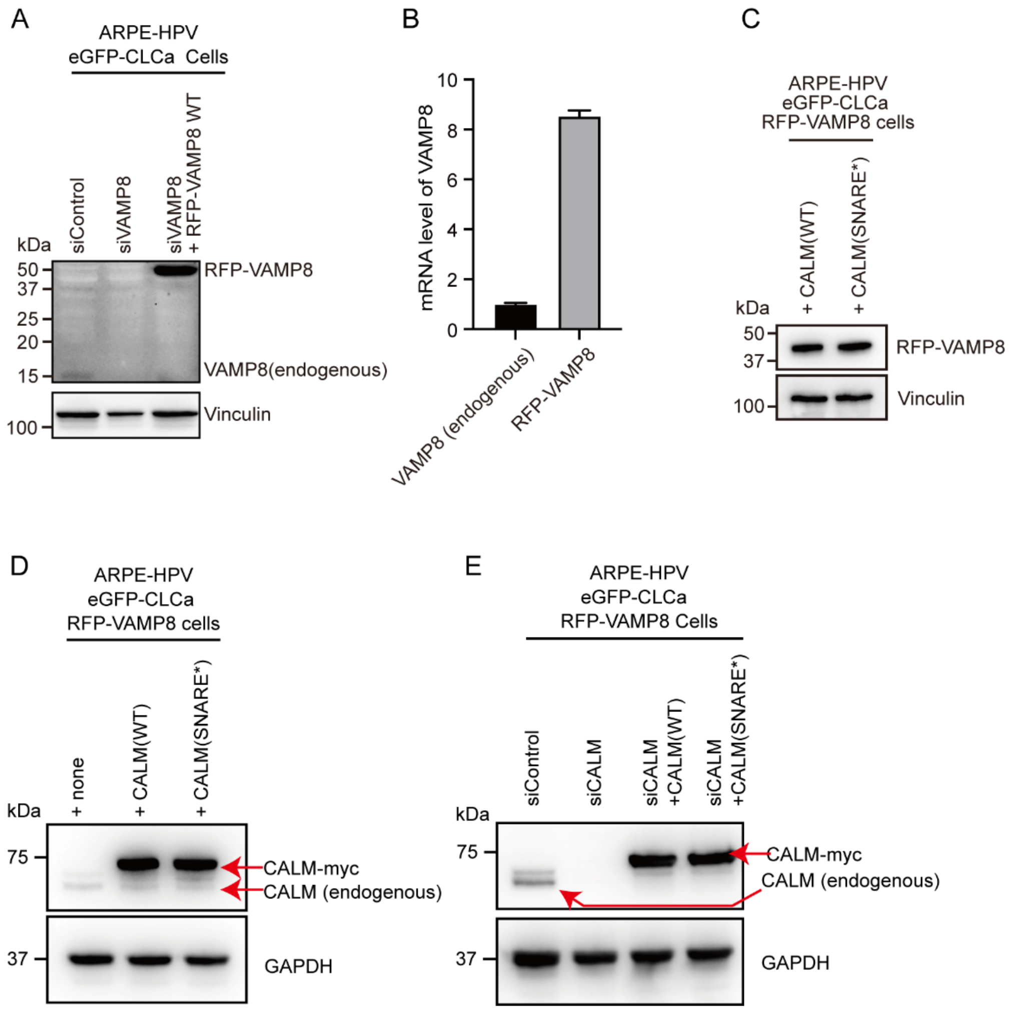

**Figure S3. Western blotting of engineered cell lines. (A)** Western blotting and RT-qPCR **(B)** indicate an ~8-fold expression of exogenous RFP-VAMP8 in ARPE-HPV eGFP-CLCa cells at the protein and RNA levels respectively. Cells were FACS sorted to have homogenous expression of

RFP-VAMP8. **(C)** Western blotting indicates the same expression level of RFP-VAMP8 in cells that stably express CALM(WT) and CALM(SNARE*). **(D)** Western blotting indicates the expression of exogenous, myc-tagged CALM(WT) and CALM(SNARE*) in ARPE-HPV eGFP-CLCa + RFP-VAMP8 cells. **(E)** CALM siRNA knocks down endogenous but not exogenous CALM in ARPE-HPV eGFP-CLCa + RFP-VAMP8 cells.

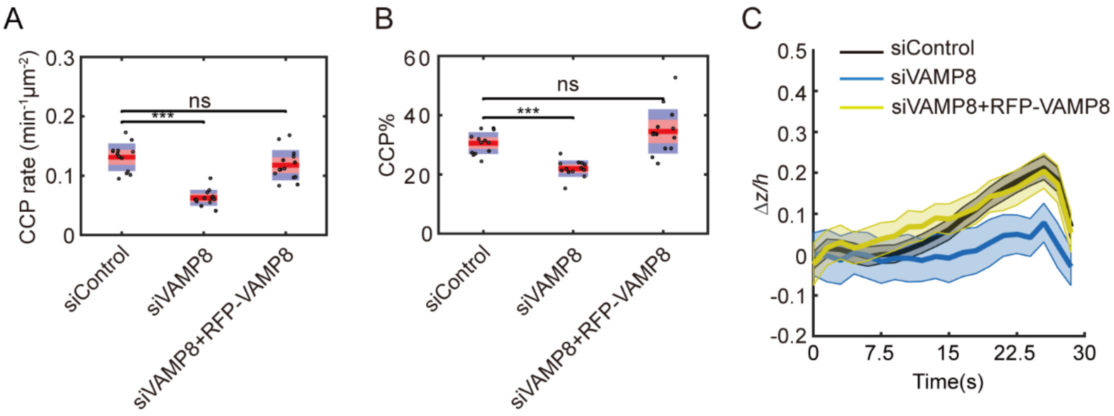

**Figure S4. Expression of siRNA-resistant RFP-VAMP8 rescues the CCP initiation, stabilization and curvature generation defects caused by VAMP8 knockdown.** Cells were transfected with control or VAMP8 siRNA. **(A-B)** TIRFM analysis revealed that stable expression of siRNA-resistant RFP-VAMP8 rescued (A) initiation rate of CCPs and (B) % of CCPs. Data presented was obtained from a single experiment (N = 13 movies for each condition). Number of dynamic tracks analyzed: 143,168 for siControl, 126,637 for siVAMP8 and 97,380 for siVAMP8+RFP-VAMP8. **(C)** Epi-TIRF microscopy analysis revealed that stable expression of siRNA-resistant RFP-VAMP8 rescued the invagination of CCPs. Data presented was obtained from a single experiment (N = 13 movies for siControl, N = 11 movies for siVAMP8, N = 13 movies for siVAMP8+RFP-VAMP8). Number of CCP tracks analyzed to obtain the $\Delta z(t)/h$ curves: 17,006 for siControl, 1,682 for siVAMP8 and 2,101 for siVAMP8+RFP-VAMP8. Statistical analysis of the data in (A-B) is the Wilcoxon Rank Sum test. ns: $P > 0.05$, $***P \leq 0.001$. Shadowed area in (C) indicates 95% confidence interval.

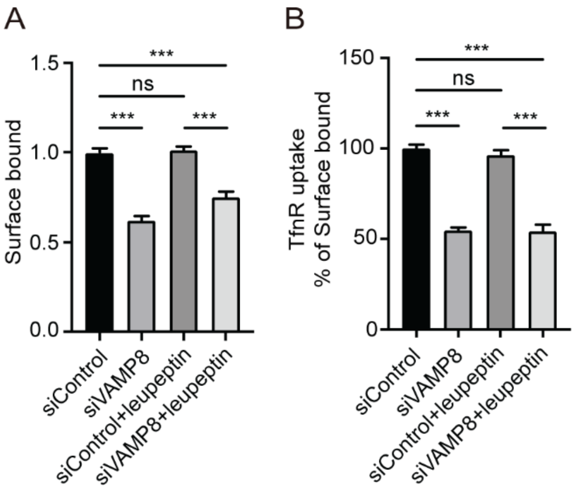

**Figure S5. TfnR uptake assay.** Measurements of the **(A)** surface bound and **(B)** uptake efficiency of TfnR. Errors indicate SEM of N = 16 samples. ***$P \leq 0.001$, ns: $P > 0.05$.

**Table S1. List of down-regulated genes after VAMP8 knockdown.**

| Gene | Protein description | Fold Change | p-value |
|---|---|---|---|
| TNFRSF10B | Tumor necrosis factor receptor superfamily member 10B | -5.2 | 6.2E-04 |
| PDGFRB | Platelet-derived growth factor receptor beta | -4.8 | 6.6E-07 |
| AXL | Tyrosine-protein kinase receptor UFO | -3.1 | 2.4E-08 |
| IL6ST | Interleukin-6 receptor subunit beta | -2.3 | 6.4E-07 |
| CXADR | Coxsackievirus and adenovirus receptor | -2.2 | 5.1E-07 |
| VLDLR | Very low-density lipoprotein receptor | -2.0 | 2.7E-06 |
| LRP8 | Low-density lipoprotein receptor-related protein 8 | -1.9 | 1.9E-05 |
| TFRC | Transferrin receptor protein 1 | -1.8 | 2.8E-06 |
| GPRC5B | G-protein coupled receptor family C group 5 member B | -1.8 | 2.7E-03 |
| EGFR | Epidermal growth factor receptor | -1.7 | 4.8E-06 |
| TGFBR1 | TGF-beta receptor type-1 | -1.6 | 4.2E-03 |
| MET | Hepatocyte growth factor receptor | -1.5 | 5.1E-05 |
| LRP1 | Prolow-density lipoprotein receptor-related protein 1 | -1.5 | 4.3E-05 |

**Table S2. List of primers.**

| Mutagenesis | Forward primer | Reverse primer |
|---|---|---|
| *VAMP8_siRNA Resistant* | *GGGGGGAAAACTTGGAACATCTCCGAAATAAAACCGAAGACCTAGAGGCCACATCTGAGCACTTCAAGA* | *TCTTGAAGTGCTCAGATGTGGCCTCTAGGTCTTCGGTTTTATTTCGGAGATGTTCCAAGTTTTCCCCCC* |
| *SNARE*: L219S* | *GTTTGCAGCATACAATGAAGGAATTATTAATTTGTCGGAAAAATATTTTGATATGAAAA* | *TTTTCATATCAAAATATTTTTCCGACAAATTAATAATTCCTTCATTGTATGCTGCAAAC* |
| *SNARE*:M244K* | *TAAGAAGTTCCTAACTAGGAAGACAAGAATCTCAGAGTTCC* | *GGAACTCTGAGATTCTTGTCTTCCTAGTTAGGAACTTCTTA* |
| **Cloning** | **Forward primer** | **Reverse primer** |
| *Backbone : pLVX-IRES-puro*<br>*insert gene: RFP-VAMP8* | *RFP* fragment: *TAGAGGATCTATTTCCGGTGCCACCATGGTAGCAGG*<br>*Linker: GCGGATCACCGCGCTTGAGAGCTACTCAGATCTCGAGCTCAAGCTT*<br>*VAMP8* fragment: *CAGTGTGCTGGAATTCGCCCTTATGGAGGAAGCCAGTGAAGG* | *RFP* fragment: *GCTTGAGCTCGAGATCTGAGTAGCTCTCAAGCGCGGTG*<br>*Linker: CACCTTCACTGGCTTCCTCCATAAGGGCGAATTCCAGCACA*<br>*VAMP8* fragment: *CGGCCGCTCTAGAACTAGTTTATGAGAAGGCACCAGTGGC* |
| *Backbone: pLVX-IRES-puro*<br>*insert gene: LAMP1-mCherry* | *TAGAGGATCTATTTCCGGTCCATGGGCGGCCCCC* | *CGGCCGCTCTAGAACTAGTCTACTTGTACAGCTCGTCCATGCC* |
